# The impact of long-term levofloxacin on the bacterial gut microbiome of young South African children

**DOI:** 10.64898/2026.02.04.703743

**Authors:** K Nel Van Zyl, AC Whitelaw, AC Hesseling, JA Seddon, A-M Demers, M Newton-Foot

## Abstract

Disruptions to gut microbial communities in early life can have lasting effects on metabolism, immune function, and resistance to infections. Antibiotics, including levofloxacin, can alter gut microbiota composition, potentially leading to long-term dysbiosis. The long-term impact of levofloxacin on the gut microbiota, especially in young children, remains poorly understood. This study investigated the effects of prolonged levofloxacin therapy over 6 months on gut microbiota in children and the stability of these changes after treatment cessation. This work used samples that were collected as part of a cluster-randomized, double-blind, placebo-controlled trial that investigated the efficacy and safety of levofloxacin for multidrug-resistant (MDR) tuberculosis (TB) preventive treatment in healthy children under the age of five years exposed to MDR-TB in the home. Levofloxacin or placebo were administered daily for 24 weeks following randomization, and stool samples were taken at baseline, and at 24- and 48-week follow-up visits. Bacterial 16S rRNA sequencing was performed on the Illumina MiSeq platform and the changes in bacterial gut microbiota composition and diversity were assessed at different time points and compared between the levofloxacin and placebo arms for different age group. Changes in the functional potential of the gut microbiome were predicted based on the observed taxonomy. Gut microbiota analysis was stratified into three age groups: 0 to <1 year, 1 to <2 years, and 2 to <5 years. The richness and evenness of microbiota were not significantly reduced following 24 weeks of levofloxacin therapy in any group. However, in infants (<1 year), the expected natural microbial diversification was significantly stunted at the end of treatment and remained impaired 24 weeks after treatment completion (48-week visit). Differential abundance testing supported this finding, revealing that a greater number of taxa were negatively impacted in the levofloxacin-treated group. Despite these shifts, beta-diversity analysis indicated no significant differences in overall microbial composition between baseline and follow-up visits after antibiotic treatment. This study showed that the natural diversification of the gut microbiota is stunted in infants and does not recover even at 24 weeks following cessation of treatment. The gut microbiota of 2 to <5-year-old children demonstrated more resilience to the influence of antibiotics.

## Introduction

The establishment of a healthy microbiome in children has been linked to long-term good health, and perturbations during this crucial phase in early life may have far-reaching implications. Antibiotic exposure is one of the most widely recognized disruptors of the human microbiota, with antibiotic-induced dysbiosis linked to increased susceptibility to secondary infections (1, 2), an increased risk of obesity (3, 4), and an increased risk of developing a pro-inflammatory conditions such as asthma, allergies, eczema, and various bowel disorders (5, 6).

While the effects of antibiotics are commonly portrayed as solely detrimental, scientific evidence suggests a more complex and context-dependent role. In rural African settings where diarrhoeal and respiratory diseases relate to a high burden of mortality and malnutrition in children, a large trial showed that twice-yearly treatment with antibiotics significantly reduced childhood mortality, although antimicrobial resistance increased and persisted in the resistome for up to 4 years (7–9).

Furthermore, the effects of antibiotics on the human microbiota are highly variable, influenced by factors such as antibiotic class, dose and duration, and the age and lifestyle of the host population (10, 11). Longitudinal studies in children are severely underrepresented in published literature, and it is therefore difficult to establish the stability of antibiotic-induced changes (11). The most microbiome studies, including those studying antibiotics, are from the Global North and represent mostly western populations (12–14). While initiatives have recently been undertaken to advance microbiome research across the African continent, particularly in humans (15), the need remains to study the microbiomes of unique populations in clinically relevant contexts across the continent (14, 16).

Levofloxacin is widely used for the treatment and prevention of multidrug-resistant (MDR) tuberculosis (TB), but its impact on the developing gut microbiota in children, outside of the field of oncology conditions remains largely unexplored. We therefore investigate the effects of a 24-week course of levofloxacin on the composition and diversity of the gut microbiota in young South African children who were enrolled in the tuberculosis child multidrug-resistant preventive therapy (TB-CHAMP) trial (17).

## Materials and Methods

### Study design

The TB-CHAMP trial assessed the efficacy and safety of levofloxacin vs. matched placebo as preventive therapy in children with household exposure to multidrug-resistant (MDR) tuberculosis (TB) (17). For the current study, participants included children aged <5 years from one of the TB-CHAMP enrolling sites and represented urban communities in the Cape Town metropolitan area (Western Cape, South Africa). These children were living in a household with an enrolled adult index patient (≥18 years of age) with bacteriologically confirmed pulmonary MDR-TB, with exclusion criteria as described previously (17). Written informed consent was obtained from the parent or legal guardian of the child enrolment in the study.

Following randomisation, child participants received either daily levofloxacin (15-20 mg/kg per day) or daily placebo treatment for six months (24 weeks). Stool samples were collected from child participants between November 2017 and March 2023 at their baseline visit (before randomisation into the treatment or placebo groups), and at the 8-, 16-, 24-, and 48-week follow-up visits. The baseline data, including microbiota sequencing (hereafter referred to as sequencing Run 1), has been previously described (18), and the sequencing data were included in this analysis. Based on the findings at baseline, participants were divided into three age groups for analyses: A (0 to <1 year), B (1 to <2 years), and C (2 to <5 years). The diversity and abundance of microbiota were compared between trial arms and at the baseline (BL), 24- and 48-week study visits overall, and within these three age groups.

Ethical approval for the trial was obtained from the Stellenbosch University Human Research Ethics Committee (SU-HREC, M16/02/009), South African Health Products Regulatory Authority (20160128) and approval was obtained to recruit from local health authorities. Ethical approval for this sub-study was also obtained from the SU-HREC (S18/02/031). The research from both the parent trial and this sub-study were conducted as outlined by Stellenbosch University’s Policy for responsible research conduct in human participants and guidelines set out by the World Health Organization (WHO) and the Declaration of Helsinki.

### Data and sample collection

Participant data was collected through clinical interviews, household questionnaires, and physical examinations during study visits. One stool sample was collected from each child at each visit in sterile 25 mL faecal containers (Lasec, South Africa) and transported to the laboratory in cooler boxes with ice packs, where they were homogenised and immediately stored at -80°C. For this study, all available 24-week and 48-week follow-up stool samples from the 115 participants (Run 1) were included. Since not all participants had stool samples available from all three visits, and additional 15 participants with the complete sample set (baseline, 24-week, and 48-week) were analysed.

### DNA extraction and sequencing

DNA was extracted from 200 mg frozen stool aliquots using the QIAamp PowerFecal DNA Isolation Kit (Qiagen, Germany) for the follow-up samples from the first 115 participants, and the baseline and follow-up samples from one additional participant, according to the manufacturer’s instructions. Due to the discontinuation of this kit, the updated QIAamp PowerFecal Pro DNA Isolation Kit (Qiagen, Germany) was used for all three samples from the remaining 14 participants. Ethanol-based purification was performed where necessary (18). As with Run 1, Run 2 was performed on the Illumina MiSeq platform at the Centre for Proteomic and Genomic Research (CPGR), Cape Town. Amplification was performed using published primers targeting the V4 hypervariable region of the 16S rRNA gene (19). Sequencing libraries were spiked with 10% of a 5 pM PhiX sequencing control. The MiSeq Reagent v3 Kit (600 cycles) was used during library preparation and 2 x 300bp paired-end reads were produced. The ZymoBIOMICS Microbial Community DNA standard (Zymo Research, USA) and batched negative extraction controls were included in the sequencing run (Supplementary material).

### Sequence analysis and statistical testing

The parameters for bioinformatic analyses may be found in the Supplementary material.

#### Sequence quality analysis

Primary data analysis was performed on the Quantitative Insights Into Microbial Ecology (QIIME2 2023.9-amplicon) bioinformatics platform (20). This included the reprocessing of Run 1 data, to ensure compatibility within the bioinformatics platform. De-multiplexed paired-end FASTQ sequences were imported, whereafter adapter and primer sequences were removed with cutadapt (21), followed by trimming of poor-quality trailing bases, error correction, quality filtering, and chimera removal using the dada2 plug-in (22). Feature tables based on amplicon sequencing variants were generated using dada2. Potential contamination was investigated by manual inspection of the negative controls and using the decontam plug-in (23) in QIIME2 (23) (Supplementary material).

#### Taxonomic profiling and differential abundance analysis

Taxonomy was assigned to the generated features using a Naïve Bayes classifier trained on the SILVA release 138 99% OTU V4 region database (https://www.arb-silva.de/silva-license-information/) (24, 25).The pipeline performance was evaluated by comparing the genus-level distribution of the microbial communities in the positive control (ZymoBIOMICS Microbial Community DNA standard, Zymo Research, USA) to their expected distribution (Supplementary Figure S1). Prior to further analysis, the baseline and follow-up feature-, sequence-, and taxonomy tables generated from Runs 1 and 2 were merged. Taxonomic features appearing in less than five samples overall, as well as non-bacterial features and features unassigned at phylum level were filtered prior to further analysis.

#### Differential abundance analysis

The Analysis of Composition of Microbiomes with Bias-Correction (ANCOM-BC) plug-in (26) in QIIME2 was used to identify differentially abundant features at phylum and genus levels, between study visits and trial arms. Inkscape v1.0.1 was used to combine and polish plots where required. Feature volatility was also investigated over study visits using the q2-longitudinal plug-in (27). Due to the potential extraction bias introduced by the updated QIAamp PowerFecal Pro DNA Isolation Kit that could affect the abundance of certain features, the baseline and follow up samples from these 14 participants were excluded from the differential abundance analysis, although these results may be found in the Supplementary Figure S2.

#### Alpha and beta diversity testing

A depth of 40000 reads was chosen for rarefaction, based on the observed feature frequency and Shannon diversity rarefaction curves of samples, whereafter seven samples were excluded from diversity analyses and 240 were retained. The alpha diversity within groups was investigated in QIIME2 using observed features, the Shannon (H) diversity metric (28), and Faith’s PD (29). Statistical significance between groups was determined using Kruskal-Wallis pairwise tests and Benjamini-Hochberg False Discovery Rate (BH-FDR) multiple test correction, where appropriate (30, 31). The dissimilarity of bacterial communities between groups were investigated with the Bray-Curtis, unweighted UniFrac, and weighted UniFrac dissimilarity metrics (32, 33). The significance of differences were determined by PERMANOVA (34) with BH-FDR adjustment where appropriate. Pairwise differences and distances between trial arms were also investigated for participants with matched samples across study visits, using the q2-longitudinal plugin (27). For all analyses, corrected p values ≤ 0.05 were considered statistically significant. Diversity data produced in QIIME2 were exported to RStudio for the visualisation of alpha- and beta diversity plots (R 4.4.2 & RStudio 2024.12.0), as described in the Supplementary material. Inkscape v1.0.1 was used to polish visualisations where required. All samples were included in diversity analysis, as all follow-up samples were extracted with a kit matching the baseline samples, and potential influences on diversity would therefore remain balanced across visits.

#### Functional prediction based on taxonomic profiles

PICRUSt2 (35) was used to predict metabolic functions by inferencing MetaCyc pathway abundances (36) from the taxonomic profiles produced in QIIME2. Pathway abundances were normalised to copies per million (CPM) and log transformed before statistical analysis using the Microbiome Multivariable Associations with Linear Models (MaAsLin) 3 tool to perform linear mixed effects modelling (37). Both age-stratified and unstratified models were included. In the age-stratified models, visit and trial arm were included as fixed effects, and their interactions were measured, whereas participants (PID) were included as a random effect. The same was true for unstratified data, with the addition of the age at sampling for each visit as a fixed effect. Minimum abundance and prevalence filters were set to 0.0001 and 0.1, respectively. Significant pathways were visualised in R (R4.4.2 & RStudio 2024.12.0), as described in the Supplementary material. As with the differential abundance testing, the final 14 samples and their follow-ups were excluded, but these results may be found in the Supplementary Table S5.

## Results

### Participant demographics and sample characteristics

A total of 130 participants were included, median age of 2.6 years (Interquartile range (IQR) 1.3 – 3.6 years), which was comparable to the parent trial median age of 2.8 years (IQR 1.3 – 4.2 years). Half of the participants were female (49.3%) and only one child living with HIV (CLWH). Three participants developed TB two or more months after randomisation; however, no follow-up samples were collected for these participants, and the baseline samples were included in analysis. A total of 247 stool samples were collected (Table 1).

**Table 1.** Distribution of stool samples (n = 247) collected from TB-CHAMP stool microbiome sub-study participants between trial arms, age groups, and study visits.

| Age group | Visit | Levofloxacin<br>(n = 109) | Placebo<br>(n = 138) | p value* |
| --- | --- | --- | --- | --- |
| A (0 to <1 year) | BL | 13 | 14 | 0.847 |
|  | 24w | 6 | 8 | 0.593 |
|  | 48w | 4 | 7 | 0.680 |
| B (1 to <2 years) | BL | 9 | 20 | **0.041** |
|  | 24w | 3 | 13 | 0.135 |
|  | 48w | 4 | 6 | 1.000 |
| C (2 to <5 years) | BL | 37 | 37 | 1.000 |
|  | 24w | 17 | 20 | 0.622 |
|  | 48w | 16 | 13 | 0.577 |

\* p values calculated with the Chi-squared test or Fisher's exact test where n < 5 BL = Baseline; 24w = 24-week follow-up; 48w = 48-week follow-up

There was a relatively equal spread of participants and samples across trial arms, except for age group B (1 to <2 years), which had more participants in the placebo arm, although this was only significant at the baseline visit (p = 0.041). Of the 130 participants included in this study, 49 (38%) only had baseline samples available, whereas less than a third of participants had matched samples across all three visits (36, 28%) (Supplemental Table S2). Matched baseline and 24- or 48-week samples were available for 31 (24%) and 14 (11%) participants, respectively (Supplemental Table S2).

### General sequencing results and taxonomy characteristics

A total of 20,106,277 paired reads were generated during the second sequencing run (Run 2) and imported into QIIME2, of which 11,232,633 paired reads remained after quality processing. Following the merging of the feature tables from Runs 1 and 2, and excluding controls, a total of 18,579,439 reads were processed for taxonomy assignment of samples, with a median of 66,393 reads/sample (IQR 57,640.5 – 75,120.5 reads/sample). Supplementary Table S3 shows the summary statistics of reads imported and retained following quality filtering in QIIME2 for both the re-processed baseline data (Run 1) and the follow-up data (Run 2).

In this study, the relative abundance of bacteria was reported at phylum (Fig 1A) and genus levels (Fig 1B). Firmicutes, Bacteroidota, Proteobacteria, and Actinobacteriota were consistently well represented across all age groups, visits and trial arms. Firmicutes and Bacteroidota were the prevalent phyla regardless of age, visit or trial arm. Following these two phyla, Verrucomicrobiota were the next most abundant, and Fusobacteriota appeared more in age group A, whereas Cyanobacteria were more common in groups B and C. At genus level, a shift toward a more diverse microbiota can be observed with increasing age, with fewer apparent differences based on visit and trial arm (Fig 1B). *Prevotella* was the most abundant genus in the older children in groups B and C.

**Figure 1.**
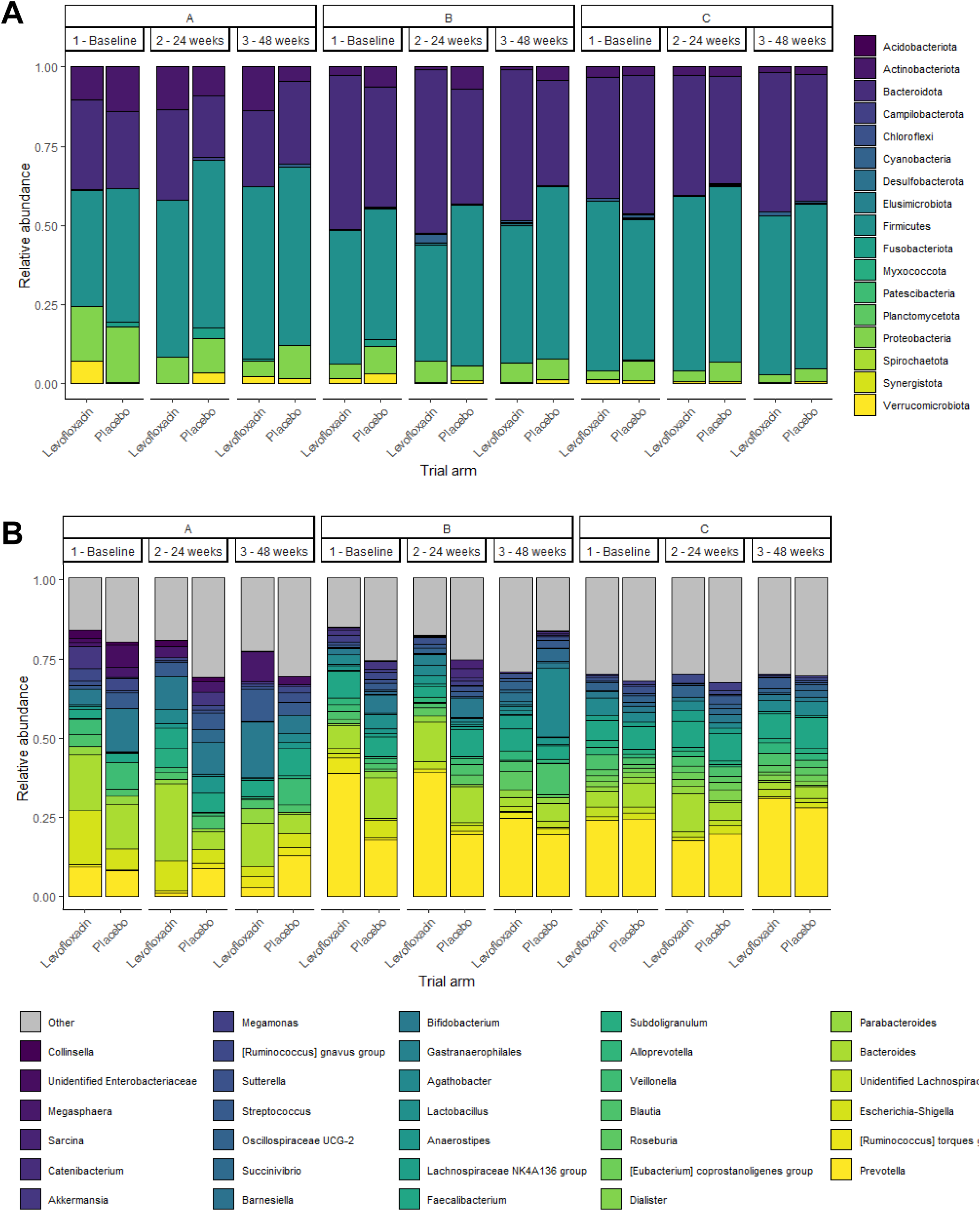
The relative abundance of bacterial phyla (A) and genera (B) identified in this study across all age groups, displayed by visit and trial arm. All detected phyla are displayed in (A), and all genera that reached the top 10 relative abundance in any of the 6 visit-trial arm comparisons per age group, are displayed in (B). Age groups = A (0 to <1 year), B (1 to <2 years), and C (2 to <5 years).

### Differential abundance at phylum and genus level over time

The ANCOM-BC plug-in in QIIME2 was used to identify differentially abundant features at phylum and genus level between the different visits in each trial arm (Fig 2). There were no differentially abundant phyla in the placebo arm in any age group at the 24-week visit, but at the 48-week visit, Actinobacteriota were depleted in both groups B and C compared to baseline. In the levofloxacin arm group A, Verrucomicrobiota were depleted at the 24-week visit and Fusobacteriota were enriched at the 48-week visit compared to baseline, and in group B, Cyanobacteria were enriched at the 48-week visit. In group C, Desulfobacterota were depleted at the 24-week visit compared to baseline.

**Figure 2.**
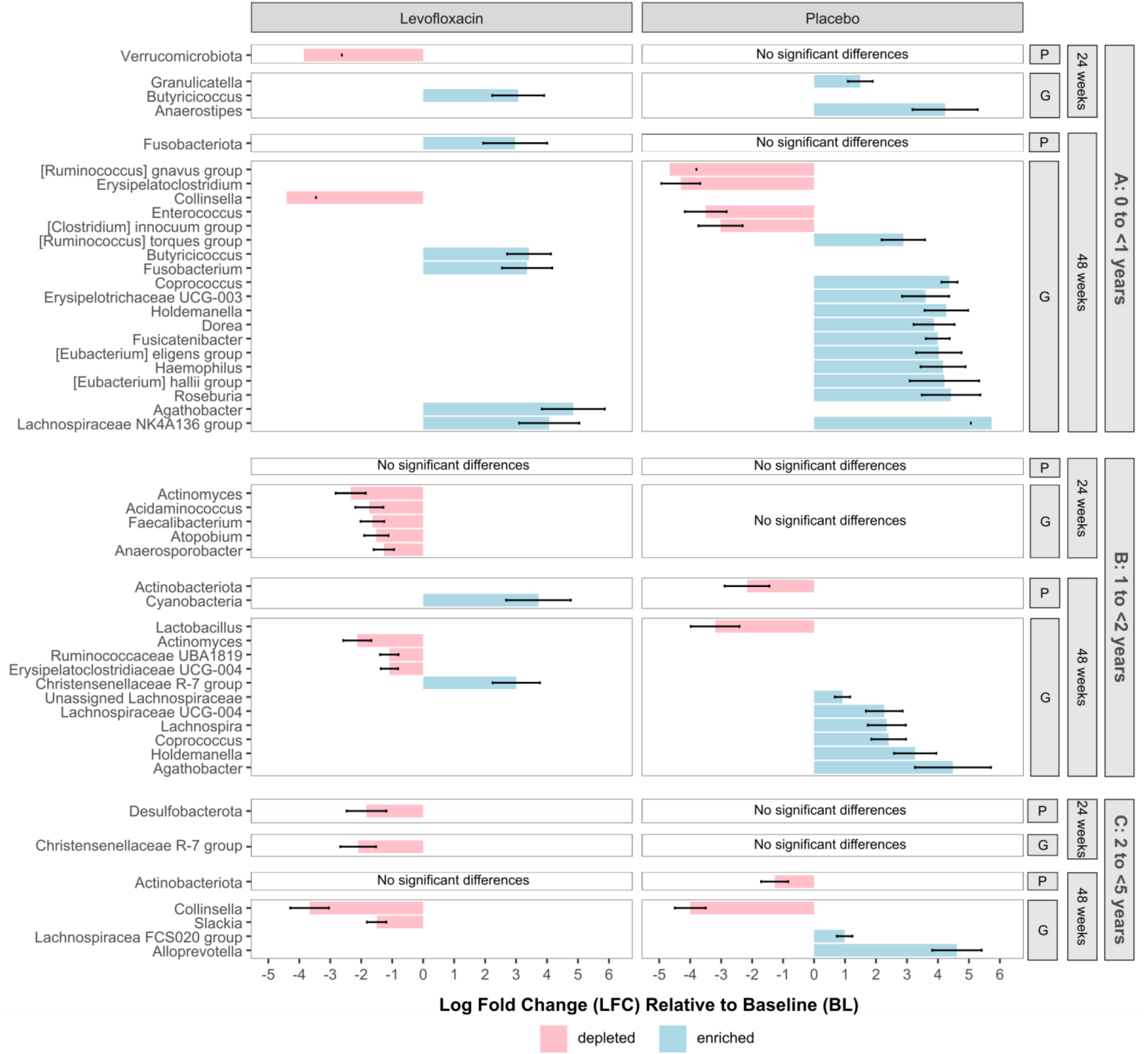
Differential abundance analysis at phylum (P) and genus (G) level, comparing the microbial composition at the 24- and 48-week follow-up visits to baseline in each trial arm, and within each age group (A (0 to <1 year), B (1 to <2 years), and C (2 to <5 years)). The log fold change (LFC) of features with significant p-values are displayed (p ≤ 0.05). Features enriched at a follow-up visit compared to baseline are shown in blue, and those depleted are shown in red.

There were more significant changes over time in the placebo arm than the levofloxacin arm at genus level, especially in group A, with a total of 11 genera significantly enriched and four depleted by the 48-week visit (Fig 2). The volatility analysis of features over time for group A (infants) identified the first three most important features as *Dorea, Butyricicoccus,* and *Anaerostipes* and their volatility patterns support what was identified by visit with ANCOM-BC: *Anaerostipes* and *Dorea* were enriched in the placebo arm at the 24- and 48-week visits, respectively, whereas *Butyricicoccus* was enriched in the levofloxacin arm, particularly at the 48-week visit.

In group B, there were five genera depleted at the 24-week visit in the levofloxacin arm, with a sustained depletion of *Actinomyces* by the 48-week visit (also observed in the full dataset, Supplementary Figure S2). The microbiota in group C were more stable between visits, regardless of trial arm, with only two and three genera significantly different at the 48-week visit, in the levofloxacin and placebo arms, respectively. *Colinsella* was depleted at this timepoint in both trial arms. When considering feature volatility, *Prevotella* was identified as the most important feature in both the older participant groups B and C, by a large margin; however, but the volatility patterns (the increase and decrease of feature abundance) were similar in the two trial arms in both groups, with the abundance of *Prevotella* rising steadily over time.

### Differences in microbial richness and evenness between study visits and trial arms

The alpha diversity of group A in the placebo arm increased across the study period and was significantly different between all visits when considering Shannon’s diversity (Fig 3A: BL-24w p = 0.048; BL-48w p = 0.002; 24w-48w p = 0.048). Conversely, those receiving levofloxacin did not diversify and the diversity remained stunted at the 48-week visit, as shown by the difference between the trial arms (p = 0.039). In this age group, the observed features also increased in the placebo group between BL and the 24- and 48-week visits (Fig 3A: p = 0.008 and p = 0.002).

**Figure 3.**
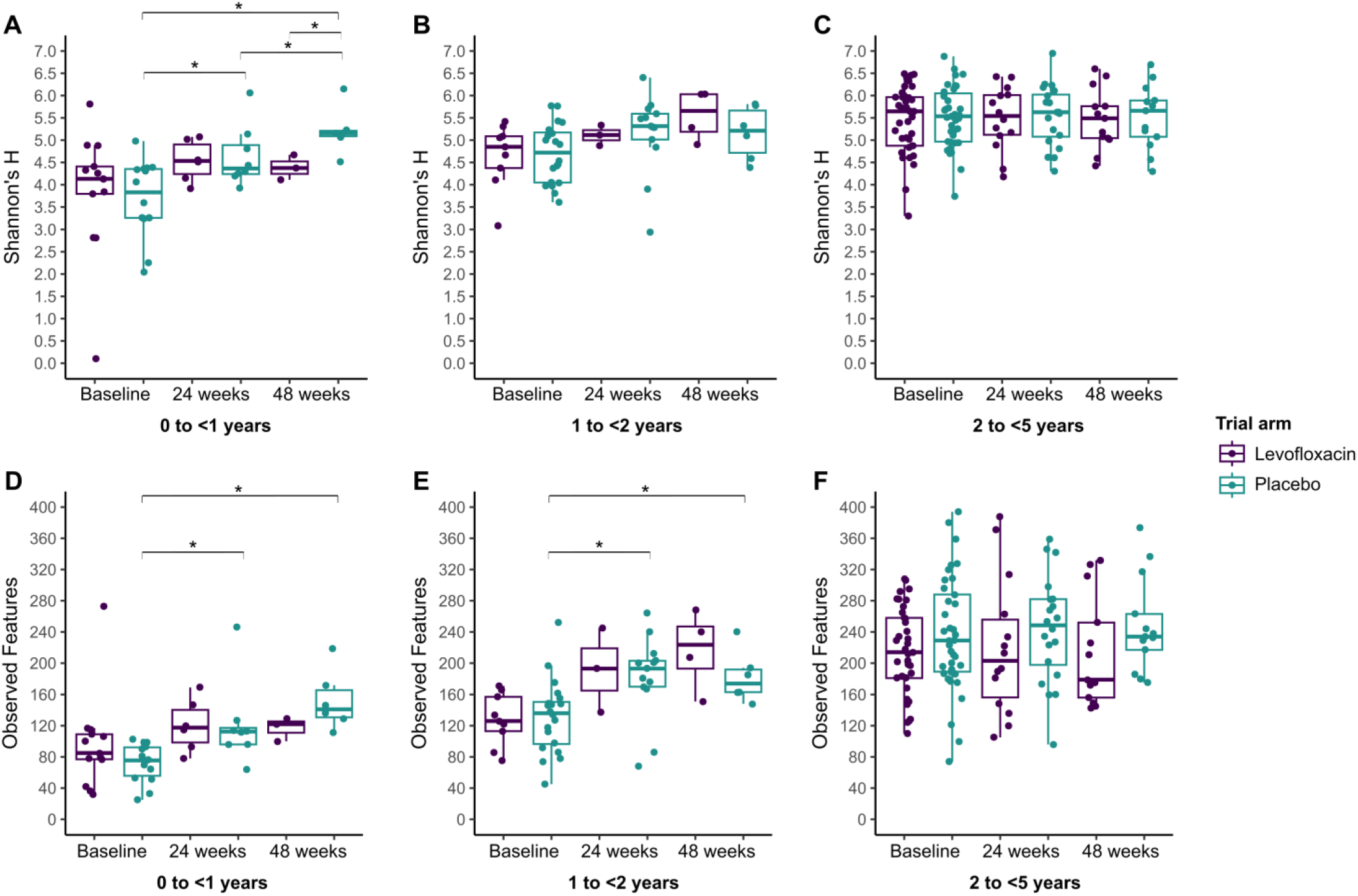
Differences in alpha diversity across the visits and trial arms, represented by Shannon’s H diversity (A, B, C) and observed features (D, E, F) in groups A (0 to <1 year), B (1 to <2 years), and C (2 to <5 years). Asterisks indicate statistically significant differences following multiple test correction (p ≤ 0.05).

When considering the number of observed features, the bacterial diversity in the placebo arm of group B had increased by the 24- and 48-week visits when compared to baseline (Fig 3B, p = 0.014 and p = 0.014). The same trend is noted in the levofloxacin arm but did not reach significance. No significant differences were noted between any groups using the Shannon’s diversity metric in group B. Diversity in group C remained stable, regardless of trial arm or visit (Fig 3C). Faith’s PD results may be viewed in the Supplementary Figure S3.

For matched samples across visits, pairwise differences in Shannon’s diversity were tested, but no significant differences were observed between visits or trial arms.

### Differences in bacterial composition as determined by dissimilarity metrics

The Bray-Curtis (BC), weighted UniFrac (WU) and unweighted UniFrac (UWU) distance measurements were used to compare bacterial composition between visits within each trial arm. In infants (group A) in the placebo arm, the bacterial composition of baseline samples was significantly different from 48-week samples by BC (p = 0.012) and UWU (p = 0.009). No other significant differences were observed between any visits in either arm. The BC dissimilarity principal coordinate (PCoA) plots when considering all data (Fig 4), and data stratified by age (Fig 5) reflect the strong influence of age and the lack of statistically significant dissimilarity in microbial composition between visits and trial arms. Furthermore, no significance was observed in the pairwise BC distances of matched samples across visits, between the two trial arms.

**Figure 4.**
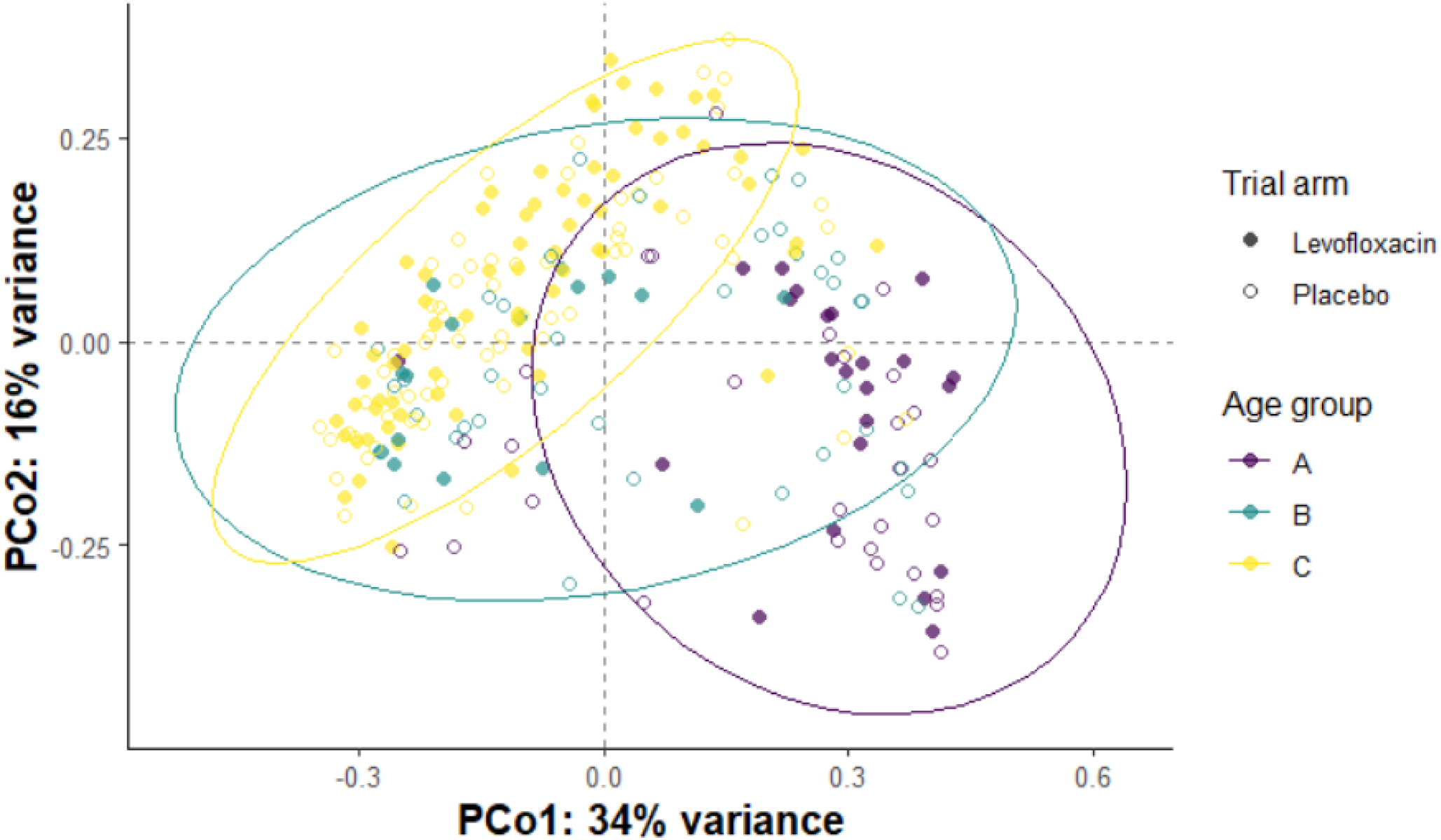
Principal Coordinate Analysis (PCoA) plots displaying the Bray-Curtis dissimilarity of gut microbiota of children, generated based on age groups A (0 to <1 year), B (1 to <2 years), and C (2 to <5 years), and trial arm (levofloxacin vs placebo). 95% confidence ellipses are shown for age groups.

**Figure 5.**
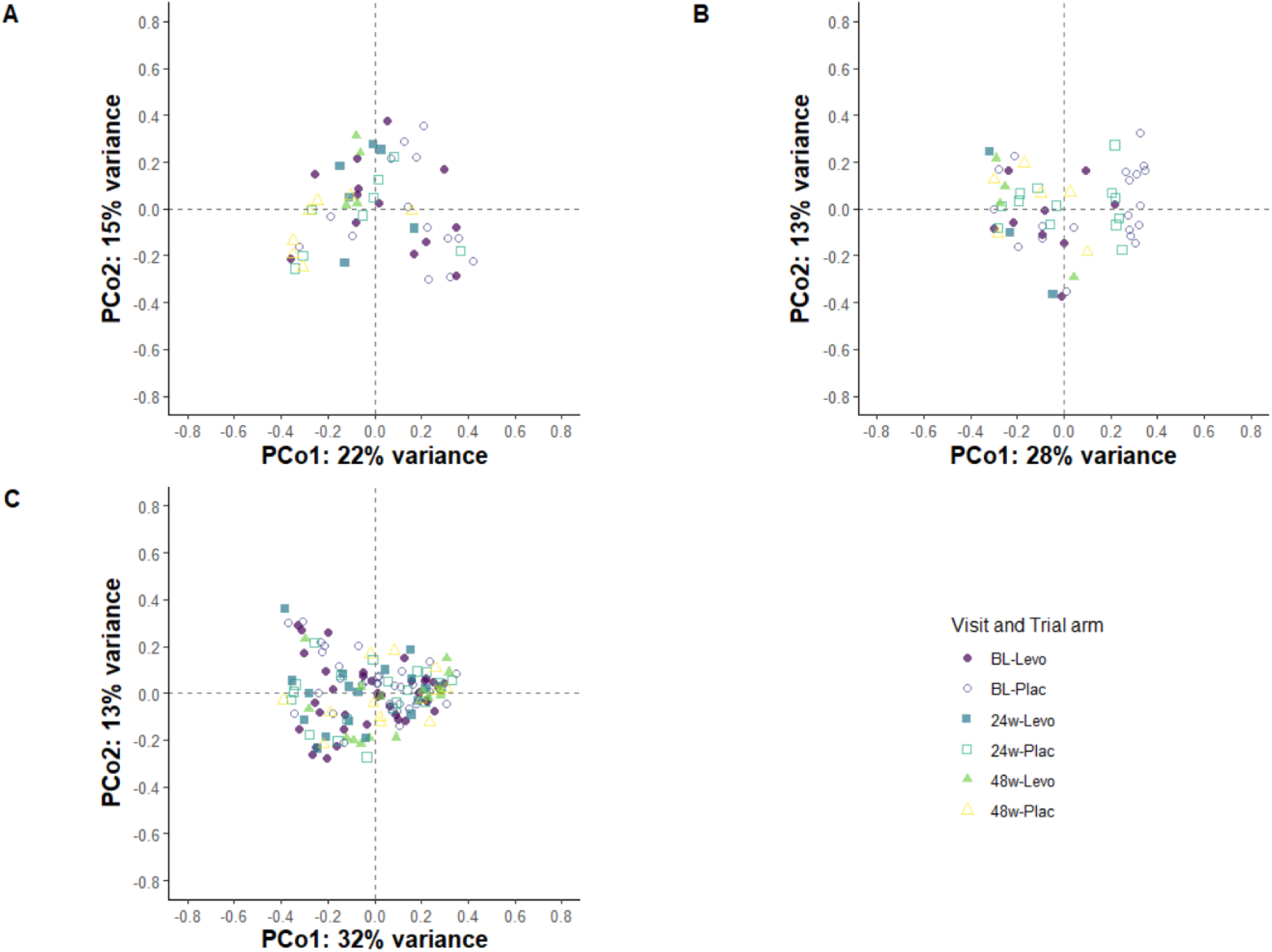
Principal Coordinate Analysis (PCoA) plots displaying the Bray-Curtis dissimilarity of gut microbiota of children, separated by age group. A) Group A (0 to <1 year), B) Group B (1 to <2 years), C) Group C (2 to <5 years). Visit (BL, 24, & 48-week) and trial arm (levofloxacin vs placebo) pairs are shown.

### Predicted functional changes in the gut microbiome

In the age-stratified analysis, no significant differences were identified for any pathways in age group A across visit or trial arm. In group B, adenosine and adenosine salvage, and the S-adenosyl-L-methionine salvage pathways were enriched at the 24-week visit, with no differences observed between trial arms. In group C, interaction effects between levofloxacin and visit were minimal, with only the 5,8-dihydroxy-2-naphthoate biosynthesis pathway showing evidence of effect association at both the 24-week and 48-week visits (Fig 6A). For individual terms, it was noted that levofloxacin was associated with the increased abundance of 79 pathways and decreased abundance in two compared to placebo, while the 24-week visit was associated with increases in two pathways and a decrease in one pathway relative to baseline (Supplementary Table S4). In the unstratified analysis, age was again shown to be a dominant driver of metabolic changes in the child gut (Supplementary Table S4); however, treatment, visit and interaction effects between treatment and visit also contributed (Supplementary Table S4, Fig 6B). At 24 weeks, 12 pathways differed between the levofloxacin and placebo groups, while at 48 weeks, only four pathways showed such differenced (Fig 6B). There were many pathways independently affected by both age and levofloxacin exposure. The pathway changes associated with age were broad, but included a shift away from specialized degradation and facultative pathways and an increase in core anabolic functions. Pathways that decreased in association with levofloxacin exposure included specialized metabolic pathways like tRNA processing, pyrimidine degradation, partial citric acid cycles, and other degradation pathways. In contrast, enrichment of core structural and biosynthetic functions, such as amino acid and nucleotide synthesis, cell membrane and wall synthesis and central metabolic pathways were observed.

**Figure 6.**
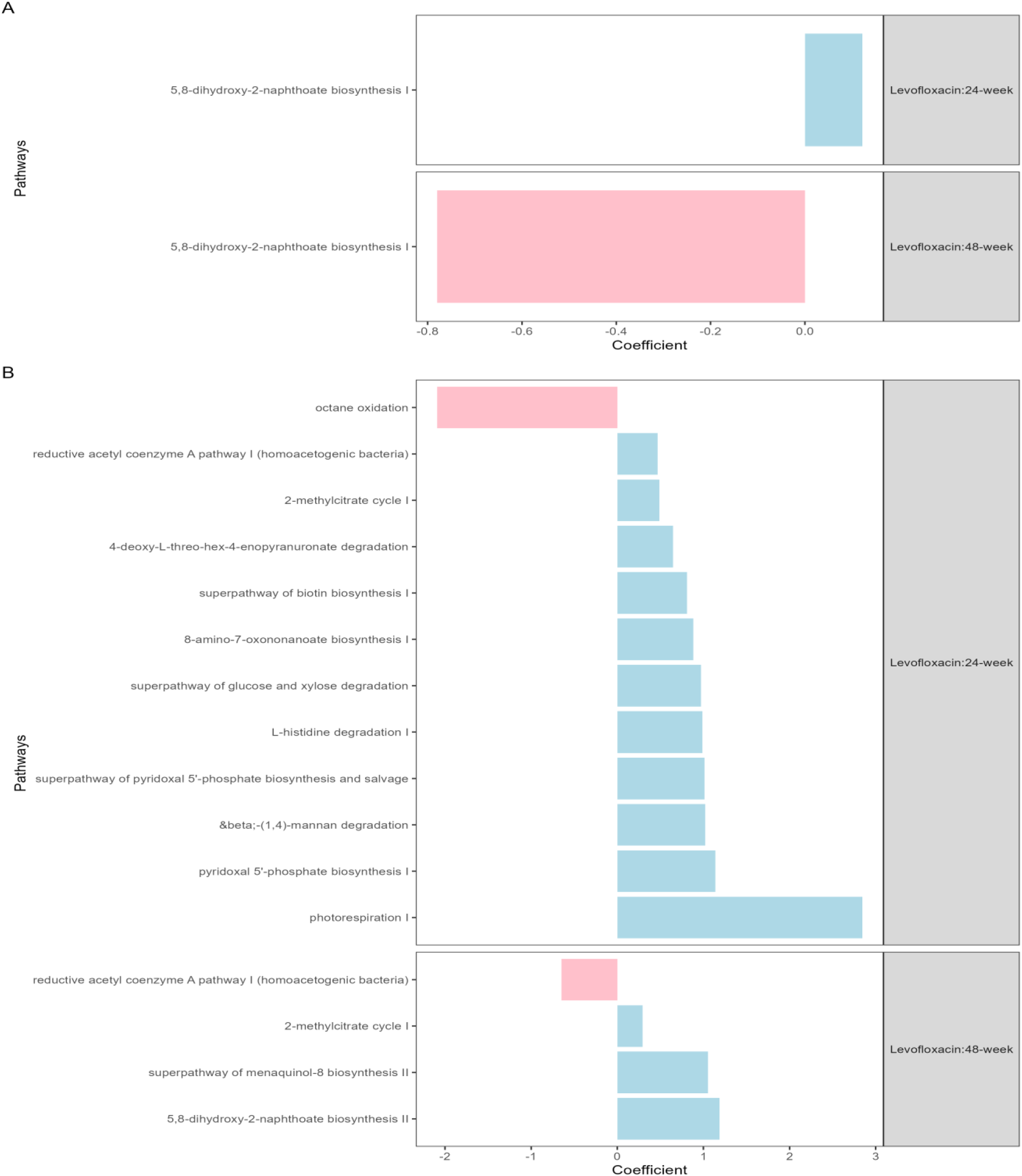
Predicted differentially abundant metabolic pathways, modelled by visit, trial arm, and the interaction of visit and trial arm. Only pathways with corrected p values (q)<0.05 were considered statistically significant and included in the visualisations. A) The stratified analysis results of age group C only (2 to <5 years) and B) The unstratified analysis of results with all samples from all age groups, with age included as a continuous variable. The results for other groups and terms are shown in Supplementary Table S4.

## Discussion

This study investigated the longitudinal effects of levofloxacin treatment on the gut microbiota and their associated functional gut pathways in young children, revealing age-dependent microbial shifts and differential responses between treatment and placebo groups. A total of 130 participants with an available stool sample at their baseline visit were included in this analysis.

A gradual diversification of bacterial microbiota was observed with increasing age that plateaued around 2 years (24 months), supporting previous findings (18). These natural changes as observed in the placebo arm complicate the interpretation of changes observed in children receiving levofloxacin, especially in the infant population. For example, in infants exposed to levofloxacin, *Butyricicoccus*, which is known to have anti-inflammatory properties and probiotic potential (38), and other butyrate producing Lachnospiraceae were elevated at the 48-week follow-up; however, *Fusobacterium* was also elevated but it, in contrast, is associated with inflammatory conditions (39, 40). This supports the holistic interpretation of findings: by combining differential abundance testing, feature volatility, diversity analysis and functional pathway analysis, we identified different butyrate producing bacteria that were enriched in each trial arm, suggesting that some microbes can adapt to antibiotic pressure and fill niches and functions left by others. In contrast, the holistic view also demonstrated how the natural increases in alpha diversity and in genera associated with the maturation of gut microbiota, including many short-chain fatty acid (SCFA) producers such as *Roseburia*, *Dorea*, *Coprococcus*, *Eubacterium*, and others (41–43), were not present in high abundance in the levofloxacin treated participants, and did not recover after a 24-week washout period. Although we did not detect statistically significant differences in predicted function in this group during stratified analysis, it has been shown that the altered microbiota play a direct role in modulating host-microbe metabolic interactions in early-life (4). Genera such as *Roseburia*, *Coprococcus*, and *Lachnospira,* which flourished in the placebo arm but not with levofloxacin exposure, have been consistently depleted in early life in children who later develop respiratory diseases like atopic wheezing and asthma (44).

Despite the small number of samples available in the levofloxacin arm at the 24-week visit, differential abundance testing identified five depleted genera compared to baseline, including *Faecalibacterium*, a microbe consistently linked to good health (45). Fluoroquinolone treatment has previously been linked to the depletion of *Faecalibacterium* in the human gut microbiota (11). However, only depletion of *Actinomyces* was sustained by the end of the study in our participants. At this final time point, there was also a trade-off in abundance between different SCFA producers in the levofloxacin arm, with enrichment of a genus level feature in the Christensenellaceae family and depletion of a genus in the Ruminococcaceae family. In the placebo arm, a significant depletion of *Lactobacillus* and increase in microbes that are often present in adult microbiota, such as *Holdemanella* and SCFA producing genera from the Lachnospiraceae family, *Coprococcus*, and *Agathobacter*, was observed at the 48-week visit, which could be attributed to the transition away from infant-like microbiota (18, 41, 46, 47). Although the gut microbiota in children between one and two years of age are still considered to be quite dynamic, we did not detect many statistically significant shifts in the microbiota in this study, even in the treatment arm. The age categories were applied at baseline during the stratification of the study data, and may therefore have masked some changes, as many of the children in this group were likely approaching the age where the microbiota start to stabilise, and would have outgrown their assigned group by the 24- and 48-week follow-up visits.

In this study, only four microbes were differentially abundant at the 48-week visit in age group C, compared to baseline. *Alloprevotella* and a genus level feature in the Lachnospiraceae FCS020 group, which are both widely detected in the human gut showed enrichment at this time point compared to baseline in the placebo arm. While there is evidence to suggest that these microbes contribute to an anti-inflammatory environment, their role in human health remains unclear (48, 49). In paediatric studies, abundance of *Alloprevotella* has contrasting associations with neurodivergence; it has been reported to be elevated in children with attention-deficit hyperactivity disorder (50), but lower in those with autism spectrum disorder (ASD) (51). Contrastingly, the Lachnospiraceae FCS020 group abundance is reportedly higher in children with ASD (52, 53). Findings from stunting studies are also inconsistent, with one study reporting a positive correlation between *Alloprevotella* abundance and stunting (54), and another a negative correlation (55). The similarity in the changes of abundance of *Collinsella* (which decreased) and *Prevotella* (which increased) in both trial arms are likely related to age-related microbiome maturation, as *Collinsella* levels have been associated with breastfeeding, whereas the establishment of *Prevotella* indicates transition to more adult-like gut microbiota (56). Overall, the stability of abundance and diversity of gut microbiota in group C, regardless of time and treatment, supports the theory that diverse gut microbiota have a greater resilience to change. This phenomenon has been explored in depth in relation to its link to human health (57) and several studies have demonstrated an association between higher diversity and positive outcomes following antibiotic treatment (58, 59). The children included in this study are exposed to MDR-TB in their households, and are likely to have a higher rate of exposure to antibiotics in the longer term compared to children exposed to M.tb, which could lead to an altered but stable, less diverse state, that is more resilient to subsequent antibiotic-induced dysbiosis (60). Evidence of microbiome resilience under antibiotic exposure was also supported by analysis of the predicted functions, which followed age-related maturation patterns typical of developing gut ecosystems, with increasing representation of host-adapted anaerobic pathways. Levofloxacin exposure was associated with a relative reduction in specialised and energy-intensive pathways and a concurrent increase in biosynthetic functions. Together, these patterns are consistent with the selective retention of functionally redundant taxa, indicating adaptation rather than functional loss.

To the best of our knowledge, this is the first study to evaluate the impact of quinolones on the gut microbiota in healthy children. Recent systematic reviews show that the majority of quinolone-related studies focused on short-term ciprofloxacin treatment in adults and do not show consistent evidence that quinolones significantly reduce overall alpha diversity, similar to our findings, although it is commonly reported that gram-negative bacilli are affected and contribute to dissimilarity of microbial composition before and after treatment in these studies (11, 61). In children, work to date comes from paediatric oncology or haematopoietic stem cell transplantation populations. Some similarities could be noted with our findings, including the depletion of *Actinomyces* and *Blautia* following prophylaxis. In contrast, while *Enterococcus* was also reported in the oncology population receiving prophylaxis, depletion was observed in the placebo arm of the current study, although only in the infant age group (62). The disruption of the early-life microbiota, including reduced microbial diversity, as observed in the present study, have been associated with adverse outcomes in children, such as increased infection risk (63), the development of atopic eczema (64), and asthma (63, 65).

In 2024, the WHO considered the results from TB-CHAMP, and another MDR-TB prevention trial (VQUIN) (66), and recommended that levofloxacin should now be used as TB preventive treatment for individuals exposed to MDR-TB (67). Countries are exploring how to scale up this intervention but there have been concerns that prolonged use of a broad-spectrum antibiotic may cause substantial disruption to the microbiome of patients. The results from this sub-study are therefore reassuring. TB-CHAMP showed a 56% reduction in the risk of developing MDR-TB in the levofloxacin group, with excellent safety and tolerance of the drug during the study period. Given the complications and mortality associated with MDR-TB disease, and the additional burden, for the child and family, of taking second-line TB treatment, the benefits of preventive levofloxacin treatment in this study population may outweigh the risks.

Our work is associated with some important limitations. While the total number of samples included in the study was substantial compared to many other longitudinal microbiome studies, the loss of retention and/or sample collections at follow-up visits were limitations in this study. Despite concerted efforts by the study team and the provision of discreet at-home collection kits, only half of the included participants provided a 24-week follow-up sample, and just over a third provided a 48-week follow-up sample. The loss of retention at 48-weeks was partly due to the lockdown imposed in South Africa during the COVID-19 pandemic in 2020 and challenges in obtaining a stool sample at scheduled study visits in children. The power of the observations was further diminished by the stratification into age groups and the exclusion of some samples from differential abundance analysis due to the necessitated changes in DNA extraction kit. To overcome this, we also included unstratified analyses with age included as a continuous variable. Further, the differential abundance results from the filtered subset, and the full dataset (Supplementary Material) showed good concordance, but with fewer differentially abundant features detected in the full set. This agreement supports our decision to retain all samples for diversity analyses, as potential extraction-related biases were minimized by processing each follow-up with the same kit used for its paired baseline. The replacement kit was also an upgraded version from the same manufacturer, with highly similar procedures. We were limited to basic taxonomy and phylogenetic inferences and functional predictions and cannot comment on the effect of levofloxacin on the gut resistome in these participants. Parallel studies are investigating the influence of antibiotic treatment in this trial on the resistance profiles of potential pathogens.

To gain a better understanding of the impact of age and antibiotic use in children, future studies could be conducted to include multi-omics techniques (e.g., metagenomics, metatranscriptomics, metabolomics, and host immune profiling) to better characterise the potential shifts in the microbiome and host-microbiome interactions following long-term levofloxacin treatment in children. This could include sequencing of stored samples that were collected at earlier timepoints during the treatment period, which may reveal temporal patterns of microbiota disruption and recovery, and distinguish resilient from non-recovering trajectories. In addition, future investigations could evaluate probiotic supplementation in infants and young children receiving TB preventive treatment as a potential means to mitigate dysbiosis. However, this will rely on a more comprehensive understanding of the microbiome and host changes observed, to inform the design of targeted and population-appropriate interventions.

Overall, our findings highlight the age-dependent effects of antibiotics on gut microbiota, with younger children displaying greater susceptibility to long-term microbial alterations. The gut microbiota of older children were more resilient to change and no significant changes in alpha or beta diversity were detected during and after 6 months’ levofloxacin, highlighting the potential association of gut microbial diversity in maintaining homeostasis. The expected diversification of the gut microbiota was stunted in infants receiving levofloxacin and did not show substantial recovery after a 24-week washout period. We hypothesize that the findings of our study, although conducted at a single site, may be broadly representative of many children living in urban areas of South Africa, where the environmental conditions, dietary patterns, and access healthcare are likely to be comparable and therefore shape similar gut microbiota profiles. While differences in gut microbiota between rural and urban populations are well recognised, the limited data available from African children underscore the need for multi-site validation with multi-omics studies to better understand the broader applicability and long-term clinical implications of antibiotic-associated gut microbiome alterations.

## Acknowledgements

We would like to thank the participants and staff of the TB-CHAMP clinical trial for collecting samples and data. We are also very grateful to Ms Anneen van Deventer, Dr Sue Purchase and Ms Elize Batist from the Desmond Tutu TB Centre for their valuable assistance. The Medical Research Council Clinical Trials Unit (MRC-CTU), UK, provided the data collected from participants. The authors would like to express their gratitude for the support given by the Division of Medical Microbiology Manuscript Writing Group.

## Author contributions

KN performed the analysis and wrote the main manuscript text. KN, AW, AH, JS and MNF contributed to the conceptualization of the study and the selection and interpretation of analyses. All authors reviewed and contributed to the manuscript.

## Additional information

This study was supported by grants funded through the NHLS Research Trust of South Africa (MNF received GRANT004_94632 and GRANT004_94679) and the Harry Crossley Foundation (KNVZ received a HCF research grant). KNVZ was supported by a postgraduate study bursary from the Harry Crossley foundation. JAS is supported by a Clinician Scientist Fellowship jointly funded by the UK Medical Research Council (MRC) and the UK Department for International Development (DFID) under the MRC/DFID Concordat agreement (MR/R007942/1). The views expressed in this article are those of the authors and do not necessarily represent the official views of the National Institutes of Health. The TB-CHAMP trial is funded by UNITAID through the Benefit-kids project to Stellenbosch University, the Joint Global Health Trials Scheme of the Department for International Development, UK (DFID), the Wellcome Trust and The Medical Research Council (MRC UK) (grant MR/M007340/1), and the South African Medical Research Council (SA MRC) Strategic Health Innovation Partnerships (SHIP) (grant S003805). The funders had no role in study design, data collection and analysis, decision to publish, or preparation of the manuscript.

