## Supplemental material for "The impact of long-term levofloxacin on the bacterial gut microbiome of young South African children"

### **Methods**

#### **DNA quality requirements**

Only DNA samples that had a concentration of at least 10 ng/μL and high purity (A_260/280_, 1.8 – 2.0; A_260/230_, 1.5 – 2.2) and integrity (≥10 Kb) could be approved for sequencing by the sequencing provider. Samples that failed to meet these standards after repeat extraction were subjected to ethanol-based purification, as previously described (1).

#### **Sequencing controls**

We included nine sequencing controls in the run: a positive control (ZymoBIOMICS Microbial Community DNA standard, Zymo Research, USA), three pairs of matched negative extraction controls for the different extraction kit lots, and another pair from the NaCl buffer used in the DNA purification process. One sample from each pair of negative controls was sequenced as is (unspiked) and the other was spiked with DNA extracted from *Streptococcus* *pneumoniae* ATCC 49614. Spiking was performed to the mean concentration of the extracted sample DNA (~90 to 100 ng/μL) to simulate true contamination levels in the negative controls and buffer. Genomic DNA was extracted from pellets taken from *S. pneumoniae* cultures that were grown shaking overnight in Brain Heart Infusion (BHI) broth (Sigma, Germany). DNA extraction was performed as described for the stool samples. We included the six negative controls and positive control from sequencing Run 1 (1) in our analysis. Read counts and the QIIME2 plug-in, decontam, were used to examine the negatives for potential contamination.

#### **Additional sequencing analyses**

As described in the main text methods, due to a discontinuation of the original DNA extraction kit, 14 baselines, and their matching follow-up stool samples, were extracted with the updated kit from the same manufacturer. These samples were excluded from differential abundance in the main text, and we show additional differential abundance analysis here for the dataset with these 42 samples included.

#### **Parameters for QIIME2 bioinformatic analyses**

For sequencing Run 1, no trailing bases were truncated during DADA2 denoising due to excellent quality throughout the reads. For sequencing Run 2, the forward reads were truncated at position 220, and the reverse reads at position 200, to remove poor quality trailing bases with median quality below Q20. The entire forward read, and the majority of the reverse reads had median quality scores of >Q30 prior to merging.

qiime taxa filter-table was employed to exclude d__Archaea, d__Eukaryota, and Unassigned features were excluded prior to analysis for both samples and mock communities. Further, only features assigned at least to Phylum level (p__) were included.

qiime taxa filter-samples was employed to filter rare features appearing in fewer than 5 samples.

#### **Packages used in R/RStudio**

A variety of packages were employed in R4.4.2 and RStudio 2024.12.0 to generate final or near-final figures (Table S1).

#### **Supplementary Table S1. R packages utilised in this study.**

| Package | Version | Applied in | Purpose |
| --- | --- | --- | --- |
| tidyverse | 2.0.0 | All figures | Data wrangling & plot drawing devices |
| ggh4x | 0.3.0 | Figure 1 | Nested facet labelling |
| viridis | 0.6.5 | Figures 1, 3, 4, & 5 | Colour-blind friendly colour palette |
| cowplot | 1.1.3 | Figures 1, 3, 4, & 5 | Plot arrangement |
| phyloseq | 1.50.0 | Figures 4 & 5 | Data wrangling for beta diversity plots |
| qiime2R | 0.99.6 | Figures 2, 4 & 5 | Conversion of QIIME2 data to phyloseq object |
| maaslin3 | 0.99.18 | Figures 2, 6 & S2 Table S4 & S5 | Differential abundance testing of microbiota and pathways using outcomes and covariates |

### **Results**

**Participant demographics and sample characteristics**

Of the 130 participants included in the study, 49 (37.7%) only had baseline samples available, whereas 67 produced a 24-week sample (51.5%), and 50 produced a 48-week sample (38.5%) (Supplementary Table S2).

#### **Supplementary Table S2. Summary of the different visit-sample combinations collected from the TB-CHAMP stool microbiome sub-study participants (n = 130).**

|  | Levofloxacin  (n = 59) | Placebo  (n = 71) | Total  (n = 130) |
| --- | --- | --- | --- |
| Participants with BL, 24wk, & 48wk samples collected | 15 (25%) | 21 (30%) | **36 (28%)** |
| Participants with BL & 24wk samples collected | 11 (19%) | 20 (28%) | **31 (24%)** |
| Participants with BL & 48wk samples collected | 9 (15%) | 5 (7%) | **14 (11%)** |
| Participants with only BL sample collected | 24 (41%) | 25 (35%) | **49 (38%)** |

#### **DNA purification and selection for sequencing**

34 DNA samples were subjected to purification following extraction, and one sample was excluded from further analysis.

#### **Negative control sequencing**

A total of 330 reads passed through dada2 filters from the three unspiked negative controls in sequencing Run 1, the majority of which originated from the salt buffer used for purification on a limited number of samples. The decontam plug-in in QIIME2 only identified 18 contaminant reads. In sequencing Run 2, only 37 reads total from the four unspiked negatives passed filters; decontam did not identify any significance. When considering the spiked negatives from either run, at least 99.95% of bacterial reads in each sample were matched to *Streptococcus* (the spike-in control). Considering these results, decontamination was not deemed necessary and was not performed on any samples.

#### **Positive control sequencing**

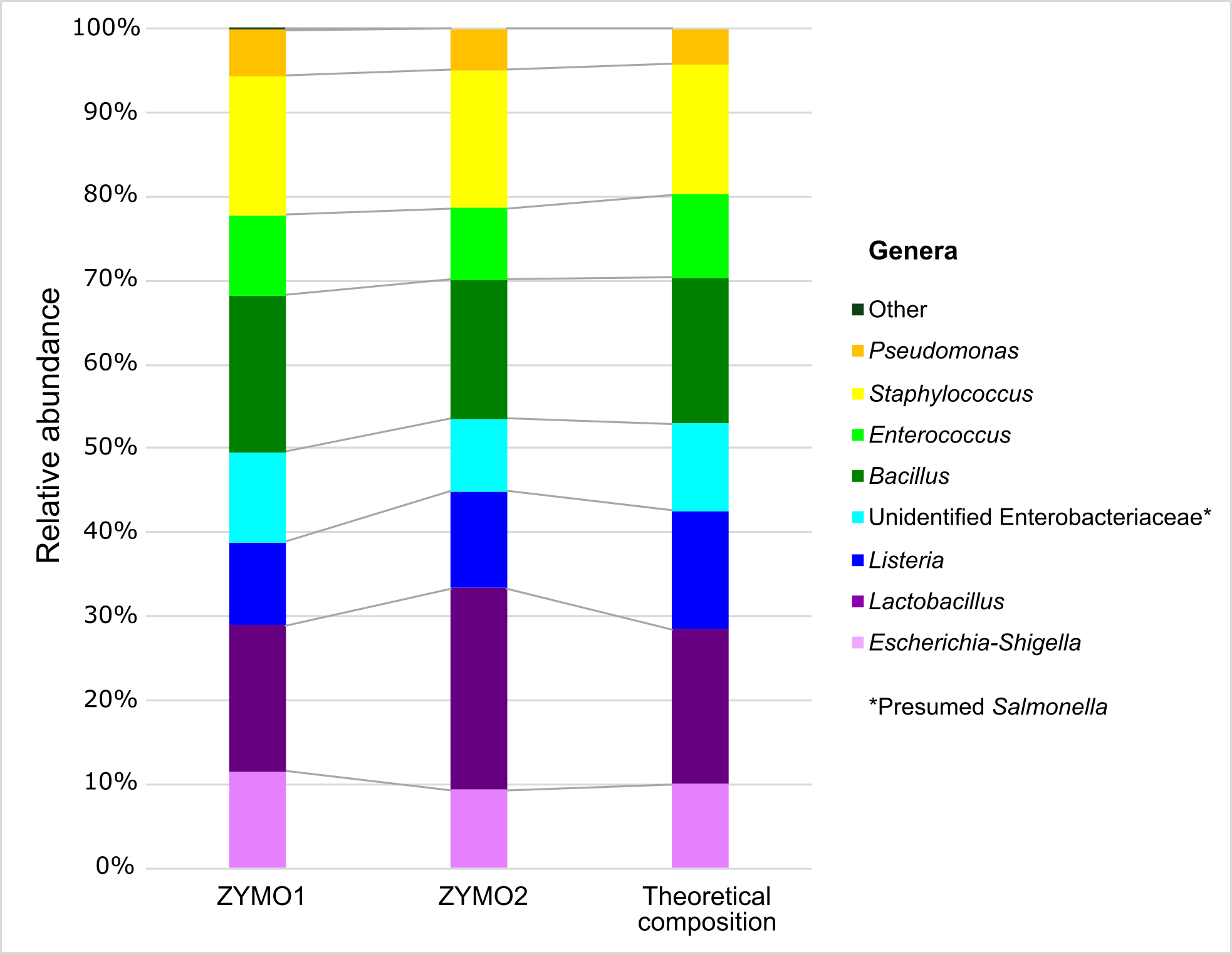
Figure S1 shows the relative abundance of detected bacterial species in the positive controls from both sequencing runs, compared to the theoretical distribution of the mock community provided by the manufacturer. *Salmonella* was not identified to genus level; however, manual Microbe Blast^TM^ of these features confirmed that the Unidentified Enterobacteriaceae in the mock samples most likely belong to *Salmonella enterica* and were therefore matched to it manually in the theoretical distribution. The inability to identify *Salmonella* was potentially due to known naming inconsistencies surrounding the genus in the SILVA138 database. A slight increase in the relative abundance of *Lactobacillus* was noted in the second sequencing run, but no substantial deviations were observed from the theoretical distribution in either experiment.

Fig S1. Comparison of the relative abundance of the positive mock community controls sequenced in run 1 (ZYMO1) and run 2 (ZYMO2) compared to the theoretical distribution. *Presumed *Salmonella*.

#### **Sequencing statistics**

#### **Supplementary Table S3. Summary read statistics for Run 1 and Run 2.**

|  | | Run 1 (baseline data) | | | | | Run 2 (follow-up and new samples) | | | | |
| --- | --- | --- | --- | --- | --- | --- | --- | --- | --- | --- | --- |
| Sample type | **Imported** | | **Passed** | **Pass %** | **Median pass** | **IQR pass** | **Imported** | **Passed** | **Pass%** | **Median pass** | **IQR pass** |
| Samples | 9160710 | | 7765590 | 85 | 67507 | 61438 – 72649 | 19352204 | 10813849 | 56 | 64793.5 | 51942 –  82111 |
| Positive mock control | 74068 | | 63838 | 86 | - | - | 194463 | 103333 | 53 | - | - |
| Unspiked negatives | 1702 | | 330 | 19 | 27 | 25 – 154 | 5123 | 37 | 0.7 | 5 | 3 – 11 |
| Spiked negatives | 244884 | | 188813 | 77 | 62926 | 62305 – 63565 | 554487 | 315414 | 57 | 81300 | 71643 – 88510 |

The poorer quality of the reverse reads in Run 2 contributed to the lower Pass rate during DADA2 denoising; however, the median reads retained per sample were similar between the two sequencing batches.

#### **Additional differential abundance results**

To determine whether the inclusion of samples from an updated sequencing kit (QIAamp PowerFecal Plus) would influence the results, we performed analysis with those 42 samples (14 baselines and their follow-ups) included. There was good agreement in the trends between the two datasets, especially at phylum level, but fewer significantly different phyla were identified overall in the all-samples dataset (Fig S2).

At genus level, this was also the case, where the main difference observed between the full set, and the dataset with samples included, shown in Fig S2, is that a more differentially abundant features were identified in the former. In the levofloxacin arm in group B, differentially abundant genera were identical in the two datasets. The microbiota in group C were mostly stable between visits, regardless of trial arm, with only *Colinsella* depleted by the 48-week visit, in both trial arms, similar to what was identified without the inclusion of all study samples.

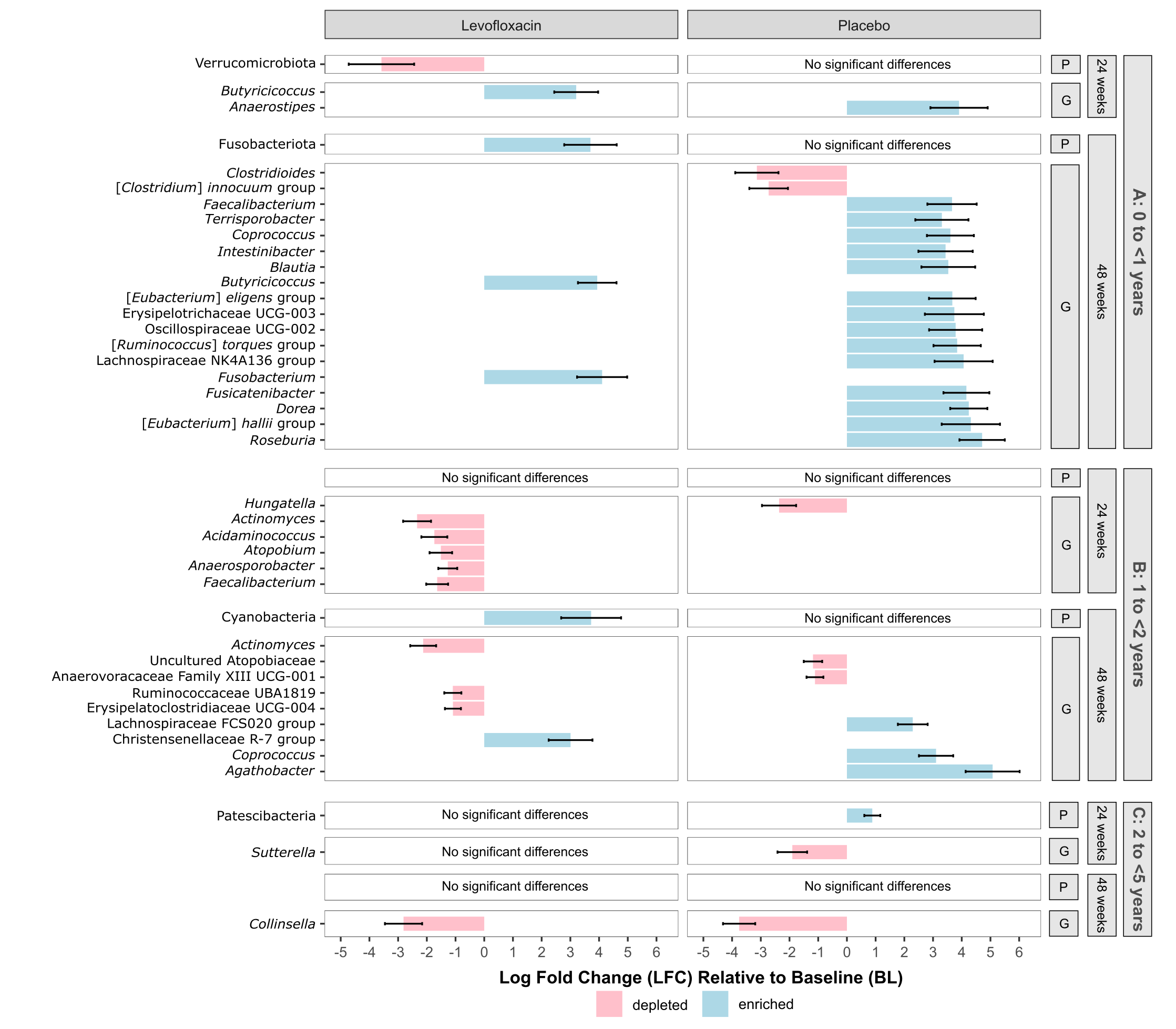

Fig S2. Differential abundance analysis at phylum (P) and genus (G) level, comparing the microbial composition at the 24- and 48-week follow-up visits to baseline in each trial arm, and within each age group. The log fold change (LFC) of features with significant p-values are displayed (p ≤ 0.05). Features enriched at a follow-up visit compared to baseline are shown in blue, and those depleted are shown in red. All samples are included.

#### **Additional alpha diversity results**

Faith’s phylogenetic diversity results are shown in Figure S3 below and follow the same trends observed with Shannon’s diversity and observed features.

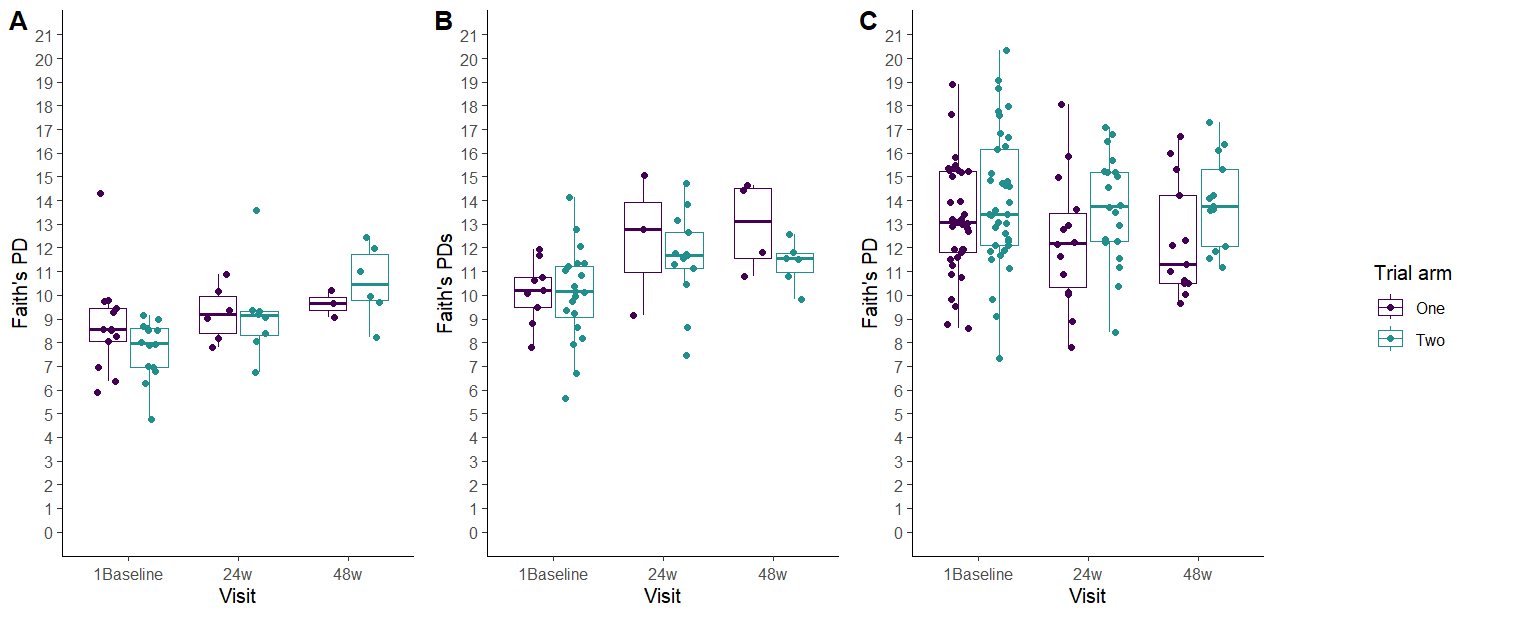

Fig S3. Differences in alpha diversity across the visits and trial arms, represented by Faith’s PD in groups A (0 to <1 years), B (1 to <2 years), and C (2 to <5 years).

#### **Additional functional pathway results**

The age-stratified and unstratified pathway analysis in MaAsLin3 was also performed with all samples included, with abundance filtering set to 0.001 and prevalence filtering to 0.1, with baseline visits and the Placebo arm set as reference levels for the association, and age included as a continuous variable in the unstratified analysis (Table S4). Age was the main driver of predicted functional changes, with 144 pathways significantly affected by increasing age. At the 24-week and 48-week visits, regardless of trial arm, 35 and 7 pathways were affected. A total of 104 pathways were associated with the levofloxacin arm, regardless of visit. In the stratified analysis, only two pathways were significantly associated with visit (24-week) for age group B, and in age group C, there were three pathways associated with visit (24-week) and 83 with the levofloxacin arm.

The age-stratified pathway analysis in MaAsLin3 was also performed with all samples included, and with the type of extraction kit used added as a fixed effect to elucidate potential kit-based biases (Table S5). Abundance filtering was set to 0.001 and prevalence filtering to 0.1, with Baseline visits, Placebo arm and the previous kit set as reference levels for the association. In age group A, three pathways were shown to be significantly correlated with kit-type and in group B, three pathways were associated with kit type and ten with trial arm, but none with visit or the interaction of visit and trial arm. In group C, 208 significant pathway-associations were identified. However, 33 of the involved pathways were associated with both the kit and trial arm, making it difficult to disentangle the individual effects of these covariates. This is most likely due to the fact that 10 of the 14 participants who had samples extracted with the new kit were randomised to the placebo arm, thereby potentially skewing the pathway abundances due to the altered extraction efficiencies of different organisms. As the sample numbers were balanced between visits, the comparison between visits, but within trial arm remain unaffected.

#### **Supplementary Table S4. Predicted functional pathways significantly associated with treatment, visit, and age (p<0.05 following correction), in the age-stratified and unstratified analyses**

| **UNSTRATIFIED ANALYSIS** | | | | | | |
| --- | --- | --- | --- | --- | --- | --- |
|  | **Pathway names** |  | **Coef** | **Std err** | **Joint p val** | **Joint q val** |
|  | pentose phosphate pathway | c | -0.44885651 | 0.069744858 | 7.26E-08 | 1.45E-05 |
|  | superpathway of methylglyoxal degradation | Age | -1.144298499 | 0.192855743 | 9.30E-08 | 1.67E-05 |
|  | catechol degradation I (meta-cleavage pathway) | Age | -1.057105384 | 0.180257676 | 1.17E-07 | 1.91E-05 |
|  | formaldehyde oxidation I | Age | -0.542543996 | 0.086871561 | 1.56E-07 | 2.34E-05 |
|  | D-galactarate degradation I | Age | -0.914937089 | 0.154644359 | 2.04E-07 | 2.66E-05 |
|  | superpathway of D-glucarate and D-galactarate degradation | Age | -0.914937089 | 0.154644359 | 2.07E-07 | 2.66E-05 |
|  | enterobactin biosynthesis | Age | -1.083625554 | 0.190021737 | 2.22E-07 | 2.66E-05 |
|  | L-arginine degradation II (AST pathway) | Age | -1.175796676 | 0.209622895 | 2.92E-07 | 2.66E-05 |
|  | superpathway of 2,3-butanediol biosynthesis (from pyruvate) | Age | -0.873185856 | 0.153028491 | 3.15E-07 | 2.66E-05 |
|  | superpathway of L-arginine and L-ornithine degradation | Age | -1.057950066 | 0.191611733 | 5.77E-07 | 3.71E-05 |
|  | superpathway of L-arginine, putrescine, and 4-aminobutanoate degradation | Age | -1.057950066 | 0.191611733 | 5.78E-07 | 3.71E-05 |
|  | superpathway of adenosylcobalamin salvage from cobinamide II | Age | 0.224263966 | 0.035349373 | 6.15E-07 | 3.76E-05 |
|  | allantoin degradation IV (anaerobic) | Age | -0.898106244 | 0.163067993 | 6.56E-07 | 3.76E-05 |
|  | adenosylcobalamin biosynthesis from adenosylcobinamide-GDP I | Age | 0.226712232 | 0.036325884 | 7.57E-07 | 3.76E-05 |
|  | mevalonate pathway I (eukaryotes and bacteria) | Age | -0.87106072 | 0.160914146 | 7.74E-07 | 3.76E-05 |
|  | superpathway of adenosylcobalamin salvage from cobinamide I | Age | 0.190005568 | 0.028348495 | 8.38E-07 | 3.97E-05 |
|  | superpathway of geranylgeranyldiphosphate biosynthesis I (via mevalonate) | Age | -0.854952854 | 0.158867522 | 9.09E-07 | 4.20E-05 |
|  | adenine and adenosine salvage III | Age | 0.154706495 | 0.017839502 | 1.08E-06 | 4.86E-05 |
|  | superpathway of sulfur oxidation (Acidianus ambivalens) | Age | 0.784996643 | 0.150416918 | 1.13E-06 | 4.94E-05 |
|  | methylphosphonate degradation I | Age | -0.834519825 | 0.15712262 | 1.41E-06 | 5.90E-05 |
|  | X3.HYDROXYPHENYLACETATE.DEGRADATION.PWY | Age | -1.096845555 | 0.203967838 | 1.72E-06 | 6.60E-05 |
|  | ketogluconate metabolism | Age | -1.078868927 | 0.207660059 | 1.76E-06 | 6.61E-05 |
|  | enterobacterial common antigen biosynthesis | Age | -1.086502608 | 0.202493502 | 1.82E-06 | 6.67E-05 |
|  | superpathway of hexitol degradation (bacteria) | Age | -0.376450818 | 0.065374332 | 1.87E-06 | 6.75E-05 |
|  | superpathway of ornithine degradation | Age | -1.021660325 | 0.196564777 | 2.05E-06 | 7.08E-05 |
|  | L-methionine biosynthesis I | Age | -0.262110317 | 0.039927118 | 2.12E-06 | 7.20E-05 |
|  | cinnamate and 3-hydroxycinnamate degradation to 2-hydroxypentadienoate | Age | -1.101837485 | 0.217002413 | 3.08E-06 | 9.54E-05 |
|  | 3-phenylpropanoate and 3-(3-hydroxyphenyl)propanoate degradation to 2-hydroxypentadienoate | Age | -1.101837485 | 0.217002413 | 3.09E-06 | 9.54E-05 |
|  | formaldehyde assimilation II (assimilatory RuMP Cycle) | Age | -0.518024755 | 0.095273639 | 3.49E-06 | 1.03E-04 |
|  | allantoin degradation to glyoxylate III | Age | -0.582722014 | 0.113928015 | 5.24E-06 | 1.37E-04 |
|  | polymyxin resistance | Age | -0.961933902 | 0.195543068 | 5.99E-06 | 1.52E-04 |
|  | sulfoquinovose degradation I | Age | -1.024212066 | 0.209816664 | 6.72E-06 | 1.66E-04 |
|  | superpathway of glycol metabolism and degradation | Age | -0.791473034 | 0.161427017 | 7.73E-06 | 1.83E-04 |
|  | CMP-legionaminate biosynthesis I | Age | 0.609386328 | 0.126132466 | 7.92E-06 | 1.85E-04 |
|  | superpathway of menaquinol-8 biosynthesis II | Age | 0.881844041 | 0.184510617 | 1.04E-05 | 2.33E-04 |
|  | NAD de novo biosynthesis I | Age | 0.176946262 | 0.030985559 | 1.08E-05 | 2.39E-04 |
|  | superpathway of S-adenosyl-L-methionine biosynthesis from L-aspartate | Age | -0.228168537 | 0.035827503 | 1.09E-05 | 2.39E-04 |
|  | 3-phenylpropanoate and 3-(3-hydroxyphenyl)propanoate degradation | Age | -0.981898528 | 0.20631463 | 1.14E-05 | 2.45E-04 |
|  | pyrimidine deoxyribonucleotides de novo biosynthesis III | Age | 0.337427307 | 0.072729713 | 1.27E-05 | 2.66E-04 |
|  | glycogen biosynthesis I (from ADP-D-Glucose) | Age | 0.140692084 | 0.020248751 | 1.36E-05 | 2.82E-04 |
|  | 5,8-dihydroxy-2-naphthoate biosynthesis I | Age | 0.829401504 | 0.208223633 | 1.40E-05 | 2.83E-04 |
|  | 2-methylcitrate cycle II | Age | -0.81409831 | 0.173408803 | 1.56E-05 | 3.12E-04 |
|  | fatty acid salvage | Age | 0.243016902 | 0.240649823 | 1.71E-05 | 3.32E-04 |
|  | superpathway of glyoxylate bypass and TCA | Age | -0.718600887 | 0.153623527 | 1.85E-05 | 3.53E-04 |
|  | 5,8-dihydroxy-2-naphthoate biosynthesis II | Age | 0.929618192 | 0.201552373 | 1.97E-05 | 3.69E-04 |
|  | superpathway of chorismate metabolism | Age | -0.659469295 | 0.137172136 | 2.72E-05 | 4.76E-04 |
|  | glyoxylate cycle | Age | -0.749899705 | 0.165330209 | 2.93E-05 | 5.03E-04 |
|  | cis-vaccenate biosynthesis | Age | 0.124740556 | 0.016911767 | 3.17E-05 | 5.33E-04 |
|  | methylerythritol phosphate pathway I | Age | 0.12012912 | 0.015410265 | 3.34E-05 | 5.52E-04 |
|  | methylerythritol phosphate pathway II | Age | 0.12012912 | 0.015410265 | 3.34E-05 | 5.52E-04 |
|  | hexitol fermentation to lactate, formate, ethanol and acetate | Age | -0.415642447 | 0.088082141 | 3.61E-05 | 5.90E-04 |
|  | superpathway of glycolysis, pyruvate dehydrogenase, TCA, and glyoxylate bypass | Age | -0.66240526 | 0.147173435 | 3.77E-05 | 6.11E-04 |
|  | inosine-5'-phosphate biosynthesis III | Age | -0.410155749 | 0.086948722 | 3.98E-05 | 6.38E-04 |
|  | D-glucarate degradation I | Age | -0.573131632 | 0.120867979 | 4.00E-05 | 6.38E-04 |
|  | 2-methylcitrate cycle I | Age | -0.816476524 | 0.184302298 | 4.14E-05 | 6.42E-04 |
|  | superpathway of Kdo2-lipid A biosynthesis | Age | -0.835424252 | 0.18811213 | 4.42E-05 | 6.75E-04 |
|  | pyrimidine deoxyribonucleosides salvage | Age | 0.122422251 | 0.017819182 | 4.99E-05 | 7.49E-04 |
|  | aerobic respiration I (cytochrome c) | Age | 0.979484548 | 0.235224323 | 5.18E-05 | 7.70E-04 |
|  | L-glutamate and L-glutamine biosynthesis | Age | 0.237824761 | 0.052327474 | 5.51E-05 | 8.13E-04 |
|  | TCA cycle IV (2-oxoglutarate decarboxylase) | Age | -0.612416786 | 0.138875831 | 5.63E-05 | 8.25E-04 |
|  | lactose degradation I | Age | -0.531082206 | 0.114200542 | 5.98E-05 | 8.60E-04 |
|  | superpathway of demethylmenaquinol-9 biosynthesis | Age | -0.618085257 | 0.140589627 | 6.42E-05 | 9.17E-04 |
|  | superpathway of demethylmenaquinol-6 biosynthesis I | Age | -0.617787887 | 0.140731596 | 6.57E-05 | 9.31E-04 |
|  | glucose and glucose-1-phosphate degradation | Age | -0.6006275 | 0.138532763 | 7.17E-05 | 0.001008714 |
|  | superpathway of (R,R)-butanediol biosynthesis (from pyruvate) | Age | -0.675667248 | 0.15129535 | 8.44E-05 | 0.001159644 |
|  | superpathway of L-aspartate and L-asparagine biosynthesis | Age | 0.113404894 | 0.016576432 | 1.02E-04 | 0.001337988 |
|  | adenosylcobalamin biosynthesis I (anaerobic) | Age | 0.49737086 | 0.220108724 | 1.12E-04 | 0.001454855 |
|  | superpathway of menaquinol-9 biosynthesis | Age | -0.570709485 | 0.134637074 | 1.20E-04 | 0.001551178 |
|  | superpathway of menaquinol-10 biosynthesis | Age | -0.570421872 | 0.134771233 | 1.21E-04 | 0.001551178 |
|  | superpathway of menaquinol-6 biosynthesis | Age | -0.570421872 | 0.134771233 | 1.22E-04 | 0.001551178 |
|  | superpathway of geranylgeranyl diphosphate biosynthesis II (via MEP) | Age | 0.106180628 | 0.014177118 | 1.37E-04 | 0.001708409 |
|  | gondoate biosynthesis (anaerobic) | Age | 0.11176328 | 0.017443947 | 1.52E-04 | 0.001875162 |
|  | partial TCA cycle (obligate autotrophs) | Age | -0.275039842 | 0.059199995 | 1.66E-04 | 0.002020522 |
|  | superpathway of polyamine biosynthesis II | Age | 0.3602929 | 0.08878287 | 1.75E-04 | 0.00210864 |
|  | photorespiration I | Age | 1.072430704 | 0.259070356 | 1.82E-04 | 0.00218998 |
|  | L-methionine biosynthesis III | Age | -0.229041464 | 0.046612503 | 1.95E-04 | 0.002330391 |
|  | glycolysis II (from fructose 6-phosphate) | Age | -0.193623481 | 0.036718432 | 2.33E-04 | 0.002723763 |
|  | fatty acid &beta;-oxidation I (generic) | Age | -0.532516382 | 0.125007793 | 2.44E-04 | 0.002811821 |
|  | superpathway of coenzyme A biosynthesis I (bacteria) | Age | 0.121835179 | 0.023605785 | 2.47E-04 | 0.002815813 |
|  | superpathway of L-serine and glycine biosynthesis I | Age | 0.106946198 | 0.016384803 | 2.65E-04 | 0.002978938 |
|  | NAD salvage pathway III (to nicotinamide riboside) | Age | -0.460224267 | 0.113673587 | 2.66E-04 | 0.002978938 |
|  | peptidoglycan biosynthesis V (&beta;-lactam resistance) | Age | -0.505345422 | 0.126483098 | 2.86E-04 | 0.003175548 |
|  | flavin biosynthesis I (bacteria and plants) | Age | 0.126337459 | 0.026300083 | 3.04E-04 | 0.003352111 |
|  | coenzyme A biosynthesis I (bacteria) | Age | 0.096692492 | 0.012311912 | 3.14E-04 | 0.003443481 |
|  | heme b biosynthesis II (oxygen-independent) | Age | -0.410949694 | 0.102182793 | 3.21E-04 | 0.003501478 |
|  | superpathway of glycerol degradation to 1,3-propanediol | Age | -0.758228049 | 0.191108122 | 4.29E-04 | 0.004491151 |
|  | mixed acid fermentation | Age | -0.159403402 | 0.027376043 | 4.55E-04 | 0.004730826 |
|  | toluene degradation I (aerobic) (via o-cresol) | Age | -0.468744806 | 0.120397301 | 4.74E-04 | 0.004884028 |
|  | PWY.5182 | Age | -0.468744806 | 0.120397301 | 4.75E-04 | 0.004884028 |
|  | superpathway of L-threonine metabolism | Age | -0.770970829 | 0.203620355 | 4.87E-04 | 0.004985517 |
|  | L-isoleucine biosynthesis IV | Age | 0.093163911 | 0.013159036 | 5.34E-04 | 0.005396267 |
|  | aspartate superpathway | Age | -0.173454925 | 0.034657524 | 7.21E-04 | 0.007054868 |
|  | GDP-D-glycero-&alpha;-D-manno-heptose biosynthesis | Age | 0.477921853 | 0.133796995 | 8.59E-04 | 0.008266779 |
|  | L-fucose degradation I | Age | -0.288958867 | 0.075273569 | 0.00101457 | 0.009517691 |
|  | (S)-propane-1,2-diol degradation | Age | -0.450840707 | 0.114711694 | 0.001018237 | 0.009517691 |
|  | superpathway of L-tryptophan biosynthesis | Age | -0.648426204 | 0.185723286 | 0.001096807 | 0.010072715 |
|  | uroporphyrinogen-III I (from glutamate) | Age | 0.119856344 | 0.02961956 | 0.001152147 | 0.010426865 |
|  | dTDP-N-acetylthomosamine biosynthesis | Age | -0.344527813 | 0.095804573 | 0.001368088 | 0.012130827 |
|  | uroporphyrinogen-III II (from glycine) | Age | 0.123812571 | 0.032512943 | 0.001403039 | 0.01233015 |
|  | superpathway of UDP-glucose-derived O-antigen building blocks biosynthesis | Age | -0.379214505 | 0.106711774 | 0.001404267 | 0.01233015 |
|  | thiamine diphosphate salvage II | Age | 0.085815316 | 0.014888659 | 0.001422955 | 0.01237352 |
|  | L-histidine biosynthesis | Age | 0.083954061 | 0.013459734 | 0.001476952 | 0.012659589 |
|  | superpathway of L-methionine biosynthesis (transsulfuration) | Age | -0.145555944 | 0.027534258 | 0.001583301 | 0.013443123 |
|  | Calvin-Benson-Bassham cycle | Age | 0.080917472 | 0.011635523 | 0.00165391 | 0.013911393 |
|  | starch degradation V | Age | 0.087535653 | 0.016342471 | 0.001674672 | 0.014020506 |
|  | tRNA charging | Age | 0.080106147 | 0.011293525 | 0.001683248 | 0.014027066 |
|  | CDP-diacylglycerol biosynthesis II | Age | 0.081335343 | 0.01294381 | 0.001767469 | 0.014461106 |
|  | CDP-diacylglycerol biosynthesis I | Age | 0.081335343 | 0.01294381 | 0.001795203 | 0.014555701 |
|  | octane oxidation | Age | 0.481515631 | 0.153651837 | 0.001855169 | 0.014974457 |
|  | inosine-5'-phosphate biosynthesis I | Age | 0.078884694 | 0.01157346 | 0.001930179 | 0.015441434 |
|  | purine ribonucleosides degradation | Age | 0.107140875 | 0.028132257 | 0.002090774 | 0.01665218 |
|  | 1,4-dihydroxy-2-naphthoate biosynthesis | Age | -0.410360808 | 0.122973214 | 0.002350107 | 0.018155332 |
|  | phosphopantothenate biosynthesis I | Age | 0.098688689 | 0.024444438 | 0.002458442 | 0.018830622 |
|  | superpathway of phylloquinol biosynthesis | Age | -0.402272132 | 0.121400351 | 0.002598651 | 0.019820218 |
|  | O-antigen building blocks biosynthesis (E. coli) | Age | 0.130618749 | 0.039517529 | 0.002717116 | 0.020497072 |
|  | L-glutamate degradation VIII (to propanoate) | Age | -0.886669811 | 0.256938579 | 0.002721556 | 0.020497072 |
|  | superpathway of pyridoxal 5'-phosphate biosynthesis and salvage | Age | -0.344109449 | 0.103409067 | 0.002884918 | 0.021547108 |
|  | superpathway of L-threonine biosynthesis | Age | 0.072889834 | 0.009642692 | 0.002965168 | 0.021971082 |
|  | myo-inositol degradation I | Age | -0.445259747 | 0.1402539 | 0.003382631 | 0.024551351 |
|  | superpathway of demethylmenaquinol-8 biosynthesis I | Age | -0.348656794 | 0.107958389 | 0.003683588 | 0.026207348 |
|  | peptidoglycan biosynthesis II (staphylococci) | Age | 0.714733077 | 0.247183543 | 0.004053072 | 0.028498159 |
|  | superpathway of glucose and xylose degradation | Age | -0.283390395 | 0.08642963 | 0.004144527 | 0.029027818 |
|  | &beta;-(1,4)-mannan degradation | Age | 0.360177356 | 0.115961932 | 0.004223977 | 0.029469609 |
|  | 5-aminoimidazole ribonucleotide biosynthesis II | Age | 0.070589777 | 0.011844652 | 0.004578204 | 0.031331662 |
|  | superpathway of pyrimidine nucleobases salvage | Age | 0.071987963 | 0.012859869 | 0.004588866 | 0.031331662 |
|  | superpathway of 5-aminoimidazole ribonucleotide biosynthesis (from PRPP) | Age | 0.070589777 | 0.011844652 | 0.004596481 | 0.031331662 |
|  | UDP-N-acetylmuramoyl-pentapeptide biosynthesis II (lysine-containing) | Age | 0.071965586 | 0.013075638 | 0.004608626 | 0.031331662 |
|  | ppGpp metabolism | Age | -0.570501465 | 0.190914212 | 0.004929777 | 0.033117169 |
|  | UDP-N-acetyl-D-glucosamine biosynthesis I | Age | 0.091249097 | 0.025723009 | 0.004930779 | 0.033117169 |
|  | adenosine ribonucleotides de novo biosynthesis | Age | 0.070236289 | 0.012426331 | 0.005060749 | 0.033863747 |
|  | pyridoxal 5'-phosphate biosynthesis I | Age | -0.334972151 | 0.107862964 | 0.005234032 | 0.034636977 |
|  | superpathway of menaquinol-13 biosynthesis | Age | -0.31573778 | 0.101919585 | 0.005658012 | 0.037221244 |
|  | superpathway of menaquinol-11 biosynthesis | Age | -0.31573778 | 0.101919585 | 0.0056659 | 0.037221244 |
|  | superpathway of menaquinol-12 biosynthesis | Age | -0.31573778 | 0.101919585 | 0.005719372 | 0.037435891 |
|  | gluconeogenesis I | Age | 0.082475512 | 0.022069991 | 0.005767296 | 0.037612799 |
|  | chorismate biosynthesis I | Age | 0.068106333 | 0.012762044 | 0.006397398 | 0.040690163 |
|  | chorismate biosynthesis from 3-dehydroquinate | Age | 0.066973124 | 0.01188216 | 0.00651723 | 0.041169853 |
|  | chondroitin sulfate degradation I (bacterial) | Age | -0.519915852 | 0.179253401 | 0.006603798 | 0.041562364 |
|  | peptidoglycan biosynthesis I (meso-diaminopimelate containing) | Age | 0.06745536 | 0.013163912 | 0.006917385 | 0.042641415 |
|  | superpathway of menaquinol-7 biosynthesis | Age | -0.300626805 | 0.098907894 | 0.006960704 | 0.042698475 |
|  | peptidoglycan biosynthesis III (mycobacteria) | Age | 0.067632218 | 0.013195268 | 0.006987682 | 0.042698475 |
|  | acetyl-CoA fermentation to butanoate | Age | 0.201205783 | 0.067604351 | 0.007243433 | 0.043752279 |
|  | superpathway of phospholipid biosynthesis III (E. coli) | Age | 0.067929291 | 0.014160846 | 0.007319705 | 0.044023587 |
|  | superpathway of fatty acid biosynthesis initiation | Age | 0.207199609 | 0.077793132 | 0.007393149 | 0.044211522 |
|  | superpathway of aromatic amino acid biosynthesis (from phosphoenolpyruvate and D-erythrose 4-phosphate) | Age | 0.065822485 | 0.012583758 | 0.00750848 | 0.044604831 |
|  | superpathway of menaquinol-8 biosynthesis I | Age | -0.297490782 | 0.099230757 | 0.007646013 | 0.045124013 |
|  | UDP-N-acetylmuramoyl-pentapeptide biosynthesis I (meso-diaminopimelate containing) | Age | 0.066571309 | 0.013256879 | 0.007720836 | 0.045416682 |
|  | palmitoleate biosynthesis I (from (5Z)-dodec-5-enoate) | Age | 0.202420255 | 0.077121151 | 0.008356534 | 0.048521809 |
|  | superpathway of L-alanine biosynthesis | 24-week visit | 0.415734447 | 0.09583027 | 4.68E-05 | 7.08E-04 |
|  | D-galactose degradation I (Leloir pathway) | 24-week visit | 0.178091623 | 0.039635181 | 2.03E-04 | 0.002409239 |
|  | superpathway of purine deoxyribonucleosides degradation | 24-week visit | 0.211439115 | 0.055680547 | 7.02E-04 | 0.006909764 |
|  | L-histidine degradation I | 24-week visit | -0.658655804 | 0.192485361 | 9.14E-04 | 0.008750173 |
|  | octane oxidation | 24-week visit | 1.3604043 | 0.403300905 | 9.81E-04 | 0.009341601 |
|  | &beta;-(1,4)-mannan degradation | 24-week visit | -0.705289207 | 0.209156077 | 0.001076588 | 0.009988958 |
|  | L-lysine biosynthesis I | 24-week visit | 0.162456716 | 0.041887586 | 0.001095102 | 0.010072715 |
|  | superpathway of geranylgeranyldiphosphate biosynthesis I (via mevalonate) | 24-week visit | 1.011774836 | 0.302806683 | 0.001127615 | 0.010303083 |
|  | lipid IVA biosynthesis (E. coli) | 24-week visit | -0.503923263 | 0.150952714 | 0.001152748 | 0.010426865 |
|  | mevalonate pathway I (eukaryotes and bacteria) | 24-week visit | 1.017594016 | 0.307219341 | 0.001238532 | 0.011146791 |
|  | superpathway of pyrimidine deoxyribonucleosides degradation | 24-week visit | 0.187326319 | 0.051418711 | 0.001303053 | 0.011669135 |
|  | superpathway of glucose and xylose degradation | 24-week visit | -0.570784435 | 0.174849514 | 0.001450475 | 0.012492131 |
|  | CMP-3-deoxy-D-manno-octulosonate biosynthesis | 24-week visit | -0.481618066 | 0.148668184 | 0.001597532 | 0.013500269 |
|  | Kdo transfer to lipid IVA (four Kdo residues) | 24-week visit | -0.492100858 | 0.153140626 | 0.001717908 | 0.014184559 |
|  | S-adenosyl-L-methionine salvage I | 24-week visit | 0.167387805 | 0.046002854 | 0.001727402 | 0.014197827 |
|  | reductive acetyl coenzyme A pathway I (homoacetogenic bacteria) | 24-week visit | -0.800342514 | 0.444197854 | 0.001783792 | 0.014528619 |
|  | pyridoxal 5'-phosphate biosynthesis I | 24-week visit | -0.741514411 | 0.238320453 | 0.002270581 | 0.017616573 |
|  | purine ribonucleosides degradation | 24-week visit | 0.200135836 | 0.059751007 | 0.002438346 | 0.018756507 |
|  | polyisoprenoid biosynthesis (E. coli) | 24-week visit | -0.235291323 | 0.073878319 | 0.002620622 | 0.019903457 |
|  | peptidoglycan biosynthesis V (&beta;-lactam resistance) | 24-week visit | 0.971980407 | 0.298472123 | 0.002966096 | 0.021971082 |
|  | superpathway of thiamine diphosphate biosynthesis I | 24-week visit | -0.243754758 | 0.079072652 | 0.003506782 | 0.025350231 |
|  | acetylene degradation (anaerobic) | 24-week visit | 0.208054219 | 0.066051204 | 0.003839615 | 0.027209869 |
|  | incomplete reductive TCA cycle | 24-week visit | -0.286227176 | 0.095464958 | 0.003985119 | 0.028130252 |
|  | pyrimidine deoxyribonucleotides de novo biosynthesis IV | 24-week visit | 0.863250465 | 0.297286679 | 0.004454443 | 0.030838454 |
|  | lactose degradation I | 24-week visit | 0.83212276 | 0.269382864 | 0.005082186 | 0.033881242 |
|  | pyrimidine deoxyribonucleotides biosynthesis from CTP | 24-week visit | 0.816232781 | 0.285504126 | 0.005120196 | 0.034008682 |
|  | phosphatidylglycerol biosynthesis II (acyl-CoA donor) | 24-week visit | 0.112532033 | 0.032281655 | 0.00651856 | 0.041169853 |
|  | phosphatidylglycerol biosynthesis I (acyl-[acp] donor) | 24-week visit | 0.112532033 | 0.032281655 | 0.006631797 | 0.041593154 |
|  | methanogenesis from acetate | 24-week visit | 0.601840358 | 0.199344455 | 0.006687215 | 0.041795095 |
|  | L-arginine biosynthesis II (acetyl cycle) | 24-week visit | 0.131816161 | 0.040953138 | 0.006782597 | 0.04217244 |
|  | L-fucose degradation I | 24-week visit | 0.382562937 | 0.137030263 | 0.006997806 | 0.042698475 |
|  | superpathway of GDP-mannose-derived O-antigen building blocks biosynthesis | 24-week visit | -0.319341611 | 0.115034497 | 0.007072403 | 0.042863048 |
|  | superpathway of N-acetylneuraminate, N-acetylglucosamine, and N-acetylmannosamine degradation to &beta;-D-fructofuranose 6-phosphate | 24-week visit | 0.245364843 | 0.086721123 | 0.007617462 | 0.045103396 |
|  | superpathway of pyridoxal 5'-phosphate biosynthesis and salvage | 24-week visit | -0.623566219 | 0.231024095 | 0.00784015 | 0.045968304 |
|  | fatty acid elongation -- saturated | 24-week visit | -0.308030192 | 0.113181784 | 0.008075492 | 0.047041702 |
|  | reductive acetyl coenzyme A pathway I (homoacetogenic bacteria) | 48-week visit | -0.212759534 | 0.532187605 | 3.38E-104 | 3.04E-101 |
|  | 2-methylcitrate cycle I | 48-week visit | -0.140107313 | 0.535794691 | 1.71E-88 | 1.03E-85 |
|  | superpathway of menaquinol-8 biosynthesis II | 48-week visit | -1.0104262 | 0.525132794 | 2.86E-07 | 2.66E-05 |
|  | 5,8-dihydroxy-2-naphthoate biosynthesis II | 48-week visit | -1.102441926 | 0.573510438 | 2.86E-07 | 2.66E-05 |
|  | pyrimidine deoxyribonucleotides de novo biosynthesis IV | 48-week visit | 0.988143492 | 0.360425786 | 0.006371662 | 0.040670183 |
|  | pyrimidine deoxyribonucleotides biosynthesis from CTP | 48-week visit | 0.940242606 | 0.345819089 | 0.00681281 | 0.04217244 |
|  | octane oxidation | 48-week visit | 1.276875716 | 0.473564821 | 0.007337265 | 0.044023587 |
|  | starch degradation V | Levofloxacin | 0.15186791 | 0.038460297 | 2.82E-08 | 8.45E-06 |
|  | L-histidine biosynthesis | Levofloxacin | 0.103422587 | 0.031323934 | 2.43E-07 | 2.66E-05 |
|  | glycogen biosynthesis I (from ADP-D-Glucose) | Levofloxacin | 0.172011655 | 0.048246097 | 3.01E-07 | 2.66E-05 |
|  | thiamine diphosphate salvage II | Levofloxacin | 0.121108181 | 0.036589167 | 3.25E-07 | 2.66E-05 |
|  | Calvin-Benson-Bassham cycle | Levofloxacin | 0.079566334 | 0.025752778 | 4.30E-07 | 3.36E-05 |
|  | superpathway of adenosylcobalamin salvage from cobinamide I | Levofloxacin | 0.279140415 | 0.072620967 | 5.15E-07 | 3.71E-05 |
|  | superpathway of geranylgeranyl diphosphate biosynthesis II (via MEP) | Levofloxacin | 0.10041751 | 0.032500665 | 5.41E-07 | 3.71E-05 |
|  | superpathway of aromatic amino acid biosynthesis (from phosphoenolpyruvate and D-erythrose 4-phosphate) | Levofloxacin | 0.085295857 | 0.028214763 | 5.42E-07 | 3.71E-05 |
|  | methylerythritol phosphate pathway II | Levofloxacin | 0.108391966 | 0.035223121 | 6.99E-07 | 3.76E-05 |
|  | L-lysine biosynthesis VI | Levofloxacin | 0.070729669 | 0.025161266 | 7.02E-07 | 3.76E-05 |
|  | chorismate biosynthesis I | Levofloxacin | 0.084351591 | 0.028624076 | 7.09E-07 | 3.76E-05 |
|  | chorismate biosynthesis from 3-dehydroquinate | Levofloxacin | 0.077970794 | 0.026825873 | 7.46E-07 | 3.76E-05 |
|  | methylerythritol phosphate pathway I | Levofloxacin | 0.108391966 | 0.035223121 | 7.67E-07 | 3.76E-05 |
|  | L-isoleucine biosynthesis IV | Levofloxacin | 0.0903836 | 0.032244134 | 1.35E-06 | 5.79E-05 |
|  | octane oxidation | Levofloxacin | 1.745990336 | 0.37840197 | 1.64E-06 | 6.59E-05 |
|  | NAD salvage pathway I (PNC VI cycle) | Levofloxacin | 0.120150731 | 0.039607358 | 1.65E-06 | 6.59E-05 |
|  | L-isoleucine biosynthesis I (from threonine) | Levofloxacin | 0.067068487 | 0.02617396 | 1.71E-06 | 6.60E-05 |
|  | L-valine biosynthesis | Levofloxacin | 0.067068487 | 0.02617396 | 1.99E-06 | 7.03E-05 |
|  | L-isoleucine biosynthesis III | Levofloxacin | 0.067996597 | 0.027119732 | 2.25E-06 | 7.48E-05 |
|  | L-lysine biosynthesis III | Levofloxacin | 0.052713363 | 0.021836932 | 2.53E-06 | 8.29E-05 |
|  | pentose phosphate pathway (non-oxidative branch) I | Levofloxacin | 0.06357774 | 0.025840752 | 2.64E-06 | 8.50E-05 |
|  | superpathway of L-isoleucine biosynthesis I | Levofloxacin | 0.048935654 | 0.021463401 | 3.13E-06 | 9.54E-05 |
|  | X1CMET2.PWY | Levofloxacin | 0.058732369 | 0.024800094 | 3.49E-06 | 1.03E-04 |
|  | CDP-diacylglycerol biosynthesis II | Levofloxacin | 0.069847919 | 0.029215371 | 4.10E-06 | 1.19E-04 |
|  | superpathway of branched chain amino acid biosynthesis | Levofloxacin | 0.062364488 | 0.026977044 | 4.46E-06 | 1.26E-04 |
|  | glycogen degradation I | Levofloxacin | 0.139165191 | 0.047751368 | 4.48E-06 | 1.26E-04 |
|  | CDP-diacylglycerol biosynthesis I | Levofloxacin | 0.069847919 | 0.029215371 | 4.56E-06 | 1.26E-04 |
|  | superpathway of phospholipid biosynthesis III (E. coli) | Levofloxacin | 0.07504571 | 0.031051666 | 4.72E-06 | 1.29E-04 |
|  | L-isoleucine biosynthesis II | Levofloxacin | 0.065705319 | 0.028086703 | 4.83E-06 | 1.30E-04 |
|  | tRNA charging | Levofloxacin | 0.054112975 | 0.024519572 | 5.20E-06 | 1.37E-04 |
|  | glycolysis III (from glucose) | Levofloxacin | 0.062295456 | 0.026965898 | 5.91E-06 | 1.52E-04 |
|  | thiazole component of thiamine diphosphate biosynthesis I | Levofloxacin | 0.11236045 | 0.042274252 | 7.29E-06 | 1.77E-04 |
|  | adenine and adenosine salvage III | Levofloxacin | 0.105550035 | 0.041185225 | 7.62E-06 | 1.83E-04 |
|  | coenzyme A biosynthesis I (bacteria) | Levofloxacin | 0.057248343 | 0.026940692 | 8.44E-06 | 1.93E-04 |
|  | NAD de novo biosynthesis I | Levofloxacin | 0.229513457 | 0.071774816 | 8.47E-06 | 1.93E-04 |
|  | L-tryptophan biosynthesis | Levofloxacin | 0.080194442 | 0.034943117 | 1.11E-05 | 2.40E-04 |
|  | 5-aminoimidazole ribonucleotide biosynthesis I | Levofloxacin | 0.039472142 | 0.021653713 | 1.19E-05 | 2.52E-04 |
|  | superpathway of L-threonine biosynthesis | Levofloxacin | 0.035124816 | 0.020998359 | 1.59E-05 | 3.15E-04 |
|  | pyruvate fermentation to isobutanol (engineered) | Levofloxacin | 0.071930683 | 0.033329692 | 1.72E-05 | 3.32E-04 |
|  | superpathway of L-serine and glycine biosynthesis I | Levofloxacin | 0.081742658 | 0.036897371 | 1.94E-05 | 3.67E-04 |
|  | S-adenosyl-L-methionine salvage I | Levofloxacin | 0.117310777 | 0.047562592 | 2.20E-05 | 4.08E-04 |
|  | 5-aminoimidazole ribonucleotide biosynthesis II | Levofloxacin | 0.042429284 | 0.024837029 | 2.26E-05 | 4.16E-04 |
|  | superpathway of 5-aminoimidazole ribonucleotide biosynthesis (from PRPP) | Levofloxacin | 0.042429284 | 0.024837029 | 2.33E-05 | 4.24E-04 |
|  | adenosine ribonucleotides de novo biosynthesis | Levofloxacin | 0.04425203 | 0.026210266 | 2.59E-05 | 4.62E-04 |
|  | inosine-5'-phosphate biosynthesis I | Levofloxacin | 0.03962413 | 0.024233495 | 2.59E-05 | 4.62E-04 |
|  | phosphatidylglycerol biosynthesis II (acyl-CoA donor) | Levofloxacin | 0.080036737 | 0.037431747 | 2.70E-05 | 4.76E-04 |
|  | phosphatidylglycerol biosynthesis I (acyl-[acp] donor) | Levofloxacin | 0.080036737 | 0.037431747 | 2.78E-05 | 4.81E-04 |
|  | D-galactose degradation I (Leloir pathway) | Levofloxacin | 0.108714264 | 0.045945298 | 3.01E-05 | 5.12E-04 |
|  | D-galactarate degradation I | Levofloxacin | -0.581740926 | 0.398367301 | 4.09E-05 | 6.40E-04 |
|  | superpathway of D-glucarate and D-galactarate degradation | Levofloxacin | -0.581740926 | 0.398367301 | 4.09E-05 | 6.40E-04 |
|  | superpathway of L-aspartate and L-asparagine biosynthesis | Levofloxacin | 0.071188812 | 0.036410544 | 4.35E-05 | 6.69E-04 |
|  | sucrose degradation III (sucrose invertase) | Levofloxacin | 0.201139697 | 0.073255321 | 5.86E-05 | 8.51E-04 |
|  | superpathway of hexuronide and hexuronate degradation | Levofloxacin | 0.261528916 | 0.090970797 | 7.82E-05 | 0.001091667 |
|  | gondoate biosynthesis (anaerobic) | Levofloxacin | 0.064523375 | 0.036476225 | 8.36E-05 | 0.001157879 |
|  | purine ribonucleosides degradation | Levofloxacin | 0.154950402 | 0.06304015 | 8.73E-05 | 0.00119064 |
|  | L-lysine biosynthesis I | Levofloxacin | 0.090112216 | 0.045210154 | 8.81E-05 | 0.001191825 |
|  | superpathway of pyrimidine deoxyribonucleosides degradation | Levofloxacin | 0.131425658 | 0.057555318 | 9.19E-05 | 0.001233818 |
|  | cis-vaccenate biosynthesis | Levofloxacin | 0.059683189 | 0.035311641 | 9.43E-05 | 0.001257658 |
|  | UMP biosynthesis I | Levofloxacin | 0.045803215 | 0.030700505 | 9.63E-05 | 0.001274084 |
|  | superpathway of purine deoxyribonucleosides degradation | Levofloxacin | 0.14743353 | 0.062803012 | 1.27E-04 | 0.001610148 |
|  | superpathway of &beta;-D-glucuronides degradation (to pyruvate) | Levofloxacin | 0.283028244 | 0.099994565 | 1.30E-04 | 0.001631919 |
|  | tRNA processing | Levofloxacin | -1.036191503 | 0.222840492 | 1.48E-04 | 0.001831451 |
|  | superpathway of coenzyme A biosynthesis I (bacteria) | Levofloxacin | 0.097888567 | 0.049497014 | 1.64E-04 | 0.002005473 |
|  | UDP-N-acetylmuramoyl-pentapeptide biosynthesis II (lysine-containing) | Levofloxacin | 0.027434528 | 0.027713942 | 2.23E-04 | 0.002618549 |
|  | peptidoglycan biosynthesis I (meso-diaminopimelate containing) | Levofloxacin | 0.025797049 | 0.027599652 | 2.42E-04 | 0.002805476 |
|  | superpathway of pyrimidine nucleobases salvage | Levofloxacin | 0.024844385 | 0.027315592 | 2.47E-04 | 0.002815813 |
|  | peptidoglycan biosynthesis III (mycobacteria) | Levofloxacin | 0.026168332 | 0.027682964 | 2.60E-04 | 0.002943241 |
|  | guanosine ribonucleotides de novo biosynthesis | Levofloxacin | 0.0278309 | 0.029587619 | 3.62E-04 | 0.003928747 |
|  | UDP-N-acetylmuramoyl-pentapeptide biosynthesis I (meso-diaminopimelate containing) | Levofloxacin | 0.022792102 | 0.027862782 | 3.66E-04 | 0.003946148 |
|  | superpathway of adenosylcobalamin salvage from cobinamide II | Levofloxacin | 0.220624234 | 0.090754053 | 3.72E-04 | 0.003983201 |
|  | L-arginine biosynthesis IV (archaea) | Levofloxacin | 0.055183585 | 0.040152017 | 3.94E-04 | 0.004191637 |
|  | pyruvate fermentation to acetate and lactate II | Levofloxacin | 0.036615522 | 0.033661757 | 4.03E-04 | 0.004269897 |
|  | L-arginine biosynthesis I (via L-ornithine) | Levofloxacin | 0.055534363 | 0.040622834 | 4.13E-04 | 0.004347511 |
|  | superpathway of adenosine nucleotides de novo biosynthesis I (from IMP and L-aspartate) | Levofloxacin | 0.013772647 | 0.025622954 | 5.33E-04 | 0.005396267 |
|  | adenosylcobalamin biosynthesis from adenosylcobinamide-GDP I | Levofloxacin | 0.216358464 | 0.093349813 | 6.17E-04 | 0.006169915 |
|  | phosphopantothenate biosynthesis I | Levofloxacin | 0.085241285 | 0.052094716 | 6.21E-04 | 0.006178744 |
|  | L-arginine biosynthesis II (acetyl cycle) | Levofloxacin | 0.065941491 | 0.046100218 | 6.66E-04 | 0.006582564 |
|  | pyrimidine deoxyribonucleosides salvage | Levofloxacin | 0.046171688 | 0.040312572 | 7.59E-04 | 0.00738837 |
|  | superpathway of heme b biosynthesis from uroporphyrinogen-III | Levofloxacin | -1.024110525 | 0.263328192 | 8.11E-04 | 0.007845649 |
|  | superpathway of histidine, purine, and pyrimidine biosynthesis | Levofloxacin | -0.670977009 | 0.161945422 | 0.001020508 | 0.009517691 |
|  | L-ornithine biosynthesis I | Levofloxacin | 0.037662929 | 0.040408262 | 0.001333445 | 0.01188218 |
|  | pyrimidine deoxyribonucleotides de novo biosynthesis IV | Levofloxacin | 0.878149525 | 0.307906471 | 0.001418793 | 0.01237352 |
|  | dTDP-N-acetylthomosamine biosynthesis | Levofloxacin | 0.560244079 | 0.211304543 | 0.001579718 | 0.013443123 |
|  | pyrimidine deoxyribonucleotides biosynthesis from CTP | Levofloxacin | 0.815289364 | 0.293869422 | 0.001716342 | 0.014184559 |
|  | superpathway of adenosine nucleotides de novo biosynthesis II (from IMP and L-aspartate) | Levofloxacin | 0.001634681 | 0.027195455 | 0.001921834 | 0.015441434 |
|  | superpathway of N-acetylneuraminate, N-acetylglucosamine, and N-acetylmannosamine degradation to &beta;-D-fructofuranose 6-phosphate | Levofloxacin | 0.184772039 | 0.094669016 | 0.002109624 | 0.016728299 |
|  | peptidoglycan recycling I | Levofloxacin | -0.848762016 | 0.231303766 | 0.002170166 | 0.016983908 |
|  | inosine-5'-phosphate biosynthesis III | Levofloxacin | -0.687414221 | 0.179700059 | 0.002268963 | 0.017616573 |
|  | pyrimidine deoxyribonucleotides de novo biosynthesis I | Levofloxacin | -0.375418955 | 0.07973634 | 0.003034944 | 0.022265073 |
|  | sucrose degradation IV (sucrose phosphorylase) | Levofloxacin | 0.286050323 | 0.133760345 | 0.003228008 | 0.023523944 |
|  | superpathway of sulfur oxidation (Acidianus ambivalens) | Levofloxacin | 0.880437598 | 0.339592416 | 0.00365145 | 0.026207348 |
|  | partial TCA cycle (obligate autotrophs) | Levofloxacin | -0.522618867 | 0.133320116 | 0.003654941 | 0.026207348 |
|  | flavin biosynthesis I (bacteria and plants) | Levofloxacin | 0.056434726 | 0.053509762 | 0.003678449 | 0.026207348 |
|  | L-fucose degradation I | Levofloxacin | 0.349686793 | 0.160991773 | 0.004337342 | 0.030143692 |
|  | uroporphyrinogen-III I (from glutamate) | Levofloxacin | 0.103093645 | 0.074079359 | 0.004768714 | 0.03226949 |
|  | D-galacturonate degradation I | Levofloxacin | 0.11987339 | 0.08258676 | 0.00596942 | 0.038650919 |
|  | taxadiene biosynthesis (engineered) | Levofloxacin | -0.697146707 | 0.205052198 | 0.006061615 | 0.039107193 |
|  | superpathway of glycolysis and the Entner-Doudoroff pathway | Levofloxacin | -0.500769068 | 0.134178947 | 0.006100951 | 0.0392204 |
|  | uroporphyrinogen-III II (from glycine) | Levofloxacin | 0.110677707 | 0.079550152 | 0.006263135 | 0.040119729 |
|  | urea cycle | Levofloxacin | -0.966060201 | 0.307626423 | 0.006817878 | 0.04217244 |
|  | L-glutamate and L-glutamine biosynthesis | Levofloxacin | 0.241663461 | 0.130811438 | 0.007069673 | 0.042863048 |
|  | L-histidine degradation I | Levofloxacin | -0.718556104 | 0.21843918 | 0.007433629 | 0.044306397 |
|  | heme b biosynthesis II (oxygen-independent) | Levofloxacin | -0.727111957 | 0.224028733 | 0.007990513 | 0.046697801 |
| **STRATIFIED ANALYSIS** | | | | | | |
| **Age group** | **Pathway names** | **Covariate** | **Coef** | **Std err** | **Joint p val** | **Joint q val** |
| B | adenine and adenosine salvage III | 24-week visit | 0.305706669 | 0.069986239 | 2.30E-05 | 0.02345904 |
| B | S-adenosyl-L-methionine salvage I | 24-week visit | 0.366169062 | 0.067578111 | 2.85E-05 | 0.02345904 |
| C | 5,8-dihydroxy-2-naphthoate biosynthesis I | 24-week visit | -0.323443421 | 0.512066024 | 2.61E-06 | 3.16E-04 |
| C | octane oxidation | 24-week visit | 1.992032437 | 0.52052429 | 5.85E-04 | 0.013238313 |
| C | D-galactose degradation I (Leloir pathway) | 24-week visit | 0.208076337 | 0.062480592 | 0.001801902 | 0.036301726 |
| C | L-lysine biosynthesis VI | Levofloxacin | 0.095501563 | 0.023903957 | 5.15E-07 | 2.37E-04 |
| C | superpathway of aromatic amino acid biosynthesis (from phosphoenolpyruvate and D-erythrose 4-phosphate) | Levofloxacin | 0.113510953 | 0.030924178 | 8.49E-07 | 2.37E-04 |
| C | chorismate biosynthesis from 3-dehydroquinate | Levofloxacin | 0.107309139 | 0.028461736 | 8.58E-07 | 2.37E-04 |
| C | L-lysine biosynthesis III | Levofloxacin | 0.084052678 | 0.022202986 | 1.06E-06 | 2.37E-04 |
| C | superpathway of adenosylcobalamin salvage from cobinamide I | Levofloxacin | 0.214721417 | 0.056128239 | 1.07E-06 | 2.37E-04 |
| C | chorismate biosynthesis I | Levofloxacin | 0.113132025 | 0.031296936 | 1.14E-06 | 2.37E-04 |
| C | pyruvate fermentation to acetate and lactate II | Levofloxacin | 0.112184247 | 0.030866673 | 1.18E-06 | 2.37E-04 |
| C | superpathway of adenosylcobalamin salvage from cobinamide II | Levofloxacin | 0.215881581 | 0.056956552 | 1.38E-06 | 2.37E-04 |
| C | adenosylcobalamin biosynthesis from adenosylcobinamide-GDP I | Levofloxacin | 0.218048012 | 0.0575693 | 1.43E-06 | 2.37E-04 |
| C | X1CMET2.PWY | Levofloxacin | 0.090679357 | 0.026884276 | 2.17E-06 | 3.16E-04 |
| C | superpathway of phospholipid biosynthesis III (E. coli) | Levofloxacin | 0.116342903 | 0.034719357 | 2.53E-06 | 3.16E-04 |
| C | tRNA charging | Levofloxacin | 0.08569949 | 0.025502537 | 2.68E-06 | 3.16E-04 |
| C | superpathway of geranylgeranyl diphosphate biosynthesis II (via MEP) | Levofloxacin | 0.101456317 | 0.030689917 | 2.91E-06 | 3.20E-04 |
| C | adenosine ribonucleotides de novo biosynthesis | Levofloxacin | 0.084525919 | 0.025800536 | 3.76E-06 | 3.51E-04 |
| C | 5-aminoimidazole ribonucleotide biosynthesis I | Levofloxacin | 0.065927637 | 0.020019004 | 3.78E-06 | 3.51E-04 |
| C | guanosine ribonucleotides de novo biosynthesis | Levofloxacin | 0.079171596 | 0.024415907 | 3.93E-06 | 3.51E-04 |
| C | CDP-diacylglycerol biosynthesis I | Levofloxacin | 0.108433795 | 0.034408888 | 4.19E-06 | 3.51E-04 |
| C | coenzyme A biosynthesis I (bacteria) | Levofloxacin | 0.083332102 | 0.026425615 | 4.75E-06 | 3.51E-04 |
| C | glycolysis III (from glucose) | Levofloxacin | 0.10047766 | 0.031356104 | 4.78E-06 | 3.51E-04 |
| C | UDP-N-acetylmuramoyl-pentapeptide biosynthesis II (lysine-containing) | Levofloxacin | 0.082823418 | 0.026529239 | 4.97E-06 | 3.51E-04 |
| C | 5-aminoimidazole ribonucleotide biosynthesis II | Levofloxacin | 0.073132197 | 0.023222399 | 5.28E-06 | 3.51E-04 |
| C | CDP-diacylglycerol biosynthesis II | Levofloxacin | 0.108433795 | 0.034408888 | 5.39E-06 | 3.51E-04 |
| C | UMP biosynthesis I | Levofloxacin | 0.093780987 | 0.030368054 | 5.43E-06 | 3.51E-04 |
| C | superpathway of 5-aminoimidazole ribonucleotide biosynthesis (from PRPP) | Levofloxacin | 0.073132197 | 0.023222399 | 5.78E-06 | 3.51E-04 |
| C | peptidoglycan biosynthesis III (mycobacteria) | Levofloxacin | 0.07847672 | 0.025700698 | 5.98E-06 | 3.51E-04 |
| C | Calvin-Benson-Bassham cycle | Levofloxacin | 0.083177873 | 0.027144632 | 6.12E-06 | 3.51E-04 |
| C | methylerythritol phosphate pathway I | Levofloxacin | 0.090159743 | 0.029850753 | 6.47E-06 | 3.51E-04 |
| C | methylerythritol phosphate pathway II | Levofloxacin | 0.090159743 | 0.029850753 | 6.63E-06 | 3.51E-04 |
| C | superpathway of pyrimidine nucleobases salvage | Levofloxacin | 0.076604393 | 0.025856708 | 6.69E-06 | 3.51E-04 |
| C | peptidoglycan biosynthesis I (meso-diaminopimelate containing) | Levofloxacin | 0.076657192 | 0.025743868 | 6.80E-06 | 3.51E-04 |
| C | pentose phosphate pathway (non-oxidative branch) I | Levofloxacin | 0.089511706 | 0.03042264 | 7.59E-06 | 3.73E-04 |
| C | superpathway of L-isoleucine biosynthesis I | Levofloxacin | 0.070215778 | 0.024008217 | 7.84E-06 | 3.73E-04 |
| C | UDP-N-acetylmuramoyl-pentapeptide biosynthesis I (meso-diaminopimelate containing) | Levofloxacin | 0.076272797 | 0.025953992 | 7.91E-06 | 3.73E-04 |
| C | superpathway of L-threonine biosynthesis | Levofloxacin | 0.062582268 | 0.021251954 | 9.19E-06 | 4.22E-04 |
| C | pyrimidine deoxyribonucleotides de novo biosynthesis IV | Levofloxacin | 1.143635002 | 0.270390285 | 9.68E-06 | 4.32E-04 |
| C | L-isoleucine biosynthesis I (from threonine) | Levofloxacin | 0.082977942 | 0.029827991 | 1.11E-05 | 4.59E-04 |
| C | L-isoleucine biosynthesis IV | Levofloxacin | 0.08676294 | 0.030587897 | 1.11E-05 | 4.59E-04 |
| C | inosine-5'-phosphate biosynthesis I | Levofloxacin | 0.065270283 | 0.0232492 | 1.11E-05 | 4.59E-04 |
| C | glycogen biosynthesis I (from ADP-D-Glucose) | Levofloxacin | 0.180234733 | 0.0561625 | 1.17E-05 | 4.71E-04 |
| C | pyrimidine deoxyribonucleotides biosynthesis from CTP | Levofloxacin | 1.073334382 | 0.258277682 | 1.22E-05 | 4.81E-04 |
| C | L-valine biosynthesis | Levofloxacin | 0.082977942 | 0.029827991 | 1.36E-05 | 5.21E-04 |
| C | thiamine diphosphate salvage II | Levofloxacin | 0.112256047 | 0.039052333 | 1.39E-05 | 5.21E-04 |
| C | S-adenosyl-L-methionine salvage I | Levofloxacin | 0.153173616 | 0.05024273 | 1.54E-05 | 5.65E-04 |
| C | L-glutamate and L-glutamine biosynthesis | Levofloxacin | 0.302858494 | 0.087814224 | 1.66E-05 | 5.95E-04 |
| C | sucrose degradation III (sucrose invertase) | Levofloxacin | 0.285565205 | 0.084381859 | 2.03E-05 | 7.00E-04 |
| C | D-galactose degradation I (Leloir pathway) | Levofloxacin | 0.155804947 | 0.052460074 | 2.03E-05 | 7.00E-04 |
| C | adenine and adenosine salvage III | Levofloxacin | 0.133193821 | 0.046701022 | 2.34E-05 | 7.90E-04 |
| C | starch degradation V | Levofloxacin | 0.134173113 | 0.046586003 | 2.52E-05 | 8.33E-04 |
| C | L-isoleucine biosynthesis II | Levofloxacin | 0.082643628 | 0.031952257 | 2.70E-05 | 8.75E-04 |
| C | purine ribonucleosides degradation | Levofloxacin | 0.237781218 | 0.074610113 | 2.86E-05 | 9.09E-04 |
| C | L-isoleucine biosynthesis III | Levofloxacin | 0.081414748 | 0.032482366 | 2.93E-05 | 9.15E-04 |
| C | superpathway of L-aspartate and L-asparagine biosynthesis | Levofloxacin | 0.075956207 | 0.03105665 | 3.15E-05 | 9.62E-04 |
| C | superpathway of branched chain amino acid biosynthesis | Levofloxacin | 0.07720625 | 0.031470758 | 3.20E-05 | 9.62E-04 |
| C | L-histidine biosynthesis | Levofloxacin | 0.090331457 | 0.036675996 | 4.67E-05 | 0.001361995 |
| C | superpathway of adenosine nucleotides de novo biosynthesis I (from IMP and L-aspartate) | Levofloxacin | 0.058907188 | 0.02630014 | 4.75E-05 | 0.001361995 |
| C | O-antigen building blocks biosynthesis (E. coli) | Levofloxacin | 0.107247492 | 0.042274058 | 4.78E-05 | 0.001361995 |
| C | pyruvate fermentation to isobutanol (engineered) | Levofloxacin | 0.096168121 | 0.039536743 | 5.83E-05 | 0.001631464 |
| C | octane oxidation | Levofloxacin | 1.906330156 | 0.463055156 | 6.00E-05 | 0.001652526 |
| C | superpathway of pyrimidine deoxyribonucleosides degradation | Levofloxacin | 0.195435893 | 0.068299911 | 6.45E-05 | 0.001745917 |
| C | cis-vaccenate biosynthesis | Levofloxacin | 0.058270566 | 0.02778001 | 6.65E-05 | 0.00177198 |
| C | superpathway of L-serine and glycine biosynthesis I | Levofloxacin | 0.107123294 | 0.04316932 | 7.66E-05 | 0.002007577 |
| C | L-lysine biosynthesis I | Levofloxacin | 0.139869893 | 0.053996945 | 7.79E-05 | 0.002009733 |
| C | superpathway of purine deoxyribonucleosides degradation | Levofloxacin | 0.219885176 | 0.076591303 | 8.73E-05 | 0.002217992 |
| C | gondoate biosynthesis (anaerobic) | Levofloxacin | 0.057180103 | 0.028585853 | 9.25E-05 | 0.00231525 |
| C | methanogenesis from acetate | Levofloxacin | 0.614932642 | 0.177535672 | 9.45E-05 | 0.002329423 |
| C | phosphatidylglycerol biosynthesis I (acyl-[acp] donor) | Levofloxacin | 0.104748903 | 0.045420248 | 1.26E-04 | 0.003069676 |
| C | phosphatidylglycerol biosynthesis II (acyl-CoA donor) | Levofloxacin | 0.104748903 | 0.045420248 | 1.35E-04 | 0.003227936 |
| C | NAD salvage pathway I (PNC VI cycle) | Levofloxacin | 0.116030981 | 0.049705774 | 2.10E-04 | 0.004947392 |
| C | pyrimidine deoxyribonucleosides salvage | Levofloxacin | 0.067469289 | 0.038501471 | 3.29E-04 | 0.007643503 |
| C | superpathway of adenosine nucleotides de novo biosynthesis II (from IMP and L-aspartate) | Levofloxacin | 0.045073997 | 0.030558127 | 3.76E-04 | 0.008623888 |
| C | glycogen degradation I | Levofloxacin | 0.128625874 | 0.063034729 | 7.67E-04 | 0.016925355 |
| C | L-arginine biosynthesis IV (archaea) | Levofloxacin | 0.07865291 | 0.046720162 | 7.68E-04 | 0.016925355 |
| C | L-arginine biosynthesis I (via L-ornithine) | Levofloxacin | 0.079181094 | 0.047279971 | 8.26E-04 | 0.017944415 |
| C | gluconeogenesis I | Levofloxacin | 0.018702261 | 0.024102451 | 0.001021447 | 0.021914689 |
| C | superpathway of histidine, purine, and pyrimidine biosynthesis | Levofloxacin | -0.801256484 | 0.199024366 | 0.001081792 | 0.022911805 |
| C | nitrate reduction VI (assimilatory) | Levofloxacin | 0.480142743 | 0.179158936 | 0.001363831 | 0.028519605 |
| C | L-tryptophan biosynthesis | Levofloxacin | 0.039171698 | 0.036716101 | 0.001702835 | 0.034729422 |
| C | superpathway of &beta;-D-glucuronides degradation (to pyruvate) | Levofloxacin | 0.268745764 | 0.117554556 | 0.00194387 | 0.038690045 |
| C | superpathway of coenzyme A biosynthesis I (bacteria) | Levofloxacin | 0.032754736 | 0.035498207 | 0.001982037 | 0.038980055 |
| C | partial TCA cycle (obligate autotrophs) | Levofloxacin | -0.807121594 | 0.219625372 | 0.002551044 | 0.049048556 |
| C | dTDP-L-rhamnose biosynthesis | Levofloxacin | 0.09763961 | 0.060680524 | 0.002553375 | 0.049048556 |

#### **Supplementary Table S5. All sample analysis, showing the predicted functional pathways significantly associated with covariates in an age-stratified analysis (p<0.05 following correction)**

| Age group | Pathway | Covariate | Coef | Std errr | Joint p val | Joint q val |
| --- | --- | --- | --- | --- | --- | --- |
| Group A | ASPASN.PWY | Kit | -0.42446 | 0.095252 | 1.32E-05 | 0.020713 |
|  | PWY.5384 | Kit | 1.331748 | 0.264897 | 2.87E-05 | 0.022563 |
|  | PWY.7234 | Kit | 1.073377 | 0.229895 | 0.000108 | 0.05652 |
| Group B | PWY.5384 | Kit | 1.24717 | 0.275318 | 3.28E-05 | 0.047879 |
|  | PWY.6471 | Kit | 1.130748 | 0.249388 | 0.000537 | 0.071805 |
|  | PWY.6545 | Kit | -1.77881 | 0.441262 | 0.000293 | 0.069633 |
|  | DENOVOPURINE2.PWY | Trial arm | 0.407175 | 0.104519 | 0.000542 | 0.071805 |
|  | PWY.6125 | Trial arm | 0.58336 | 0.143505 | 0.000234 | 0.069633 |
|  | PWY.7184 | Trial arm | 0.688753 | 0.167473 | 0.000179 | 0.069633 |
|  | PWY.7187 | Trial arm | 0.400889 | 0.104838 | 0.000743 | 0.07742 |
|  | PWY.7196 | Trial arm | 0.57621 | 0.153265 | 0.000646 | 0.07742 |
|  | PWY.7197 | Trial arm | 0.731611 | 0.184663 | 0.000289 | 0.069633 |
|  | PWY.7200 | Trial arm | 0.460354 | 0.122352 | 0.000734 | 0.07742 |
|  | PWY.7228 | Trial arm | 0.63274 | 0.156553 | 0.000243 | 0.069633 |
|  | PWY.841 | Trial arm | 0.434638 | 0.111799 | 0.000521 | 0.071805 |
|  | PWY0.162 | Trial arm | 0.509318 | 0.127635 | 0.000338 | 0.069633 |
|  | PWY0.166 | Trial arm | 0.453102 | 0.113887 | 0.000382 | 0.069633 |
| Group C | PWY.6467 | kit | new | -1.635938822 | 0.16878578 | 1.30E-16 |
|  | NAGLIPASYN.PWY | kit | new | -1.550954133 | 0.166902552 | 1.17E-15 |
|  | PWY.5154 | kit | new | -1.170470834 | 0.127327396 | 8.44E-15 |
|  | FASYN.ELONG.PWY | kit | new | -1.021960358 | 0.111419203 | 2.52E-14 |
|  | POLYISOPRENSYN.PWY | kit | new | -0.844354799 | 0.086899403 | 2.70E-14 |
|  | FASYN.INITIAL.PWY | kit | new | -1.42843011 | 0.165926807 | 4.28E-14 |
|  | PWY.5989 | kit | new | -1.43055133 | 0.165744253 | 3.88E-14 |
|  | PWY.6282 | kit | new | -1.405682587 | 0.164464118 | 5.84E-14 |
|  | PWY.1269 | kit | new | -1.391569217 | 0.163613536 | 7.72E-14 |
|  | PWY.7664 | kit | new | -1.29639232 | 0.152280859 | 1.03E-13 |
|  | PWYG.321 | kit | new | -1.299482017 | 0.152492561 | 9.62E-14 |
|  | THISYN.PWY | kit | new | -0.792140161 | 0.084752713 | 1.85E-13 |
|  | PWY0.862 | kit | new | -1.114326082 | 0.141727856 | 3.52E-12 |
|  | PWY.5695 | kit | new | -0.515303153 | 0.059693476 | 3.73E-10 |
|  | PWY.7323 | kit | new | -0.861627466 | 0.122969019 | 3.91E-10 |
|  | COLANSYN.PWY | kit | new | -0.654769495 | 0.098091146 | 5.42E-09 |
|  | P42.PWY | kit | new | -0.722533255 | 0.087904612 | 8.56E-09 |
|  | PRPP.PWY | kit | new | -1.151756728 | 0.197478845 | 2.83E-08 |
|  | METH.ACETATE.PWY | kit | new | 1.519170281 | 0.202081187 | 9.13E-08 |
|  | PWY.6895 | kit | new | -0.685703099 | 0.118255163 | 8.91E-08 |
|  | PWY.6519 | kit | new | -1.543310482 | 0.255324432 | 3.54E-07 |
|  | PWY.7196 | kit | new | -0.578486975 | 0.106980273 | 6.22E-07 |
|  | PWY.7315 | kit | new | 1.592679333 | 0.248304844 | 6.21E-07 |
|  | TEICHOICACID.PWY | kit | new | 1.345622579 | 0.201555015 | 5.82E-07 |
|  | BIOTIN.BIOSYNTHESIS.PWY | kit | new | -1.37725289 | 0.238268601 | 7.93E-07 |
|  | PWY.7228 | kit | new | -0.552205222 | 0.103945093 | 9.87E-07 |
|  | PWY.6125 | kit | new | -0.509740161 | 0.097730628 | 1.71E-06 |
|  | PWY.7200 | kit | new | -0.45834513 | 0.085369081 | 1.75E-06 |
|  | PWY.7184 | kit | new | -0.545791704 | 0.107025764 | 1.99E-06 |
|  | PWY.7197 | kit | new | -0.590827757 | 0.118359372 | 2.22E-06 |
